# Integrins and tetraspanins mediate CD36-actin coupling and regulate CD36 organization and signaling

**DOI:** 10.1101/2025.04.24.650515

**Authors:** Soma Jana, Huong-Tra Ngo, Jaime Guerrero, Jesus Vega-Lugo, Aparajita Dasgupta, Khuloud Jaqaman

**Affiliations:** Department of Biophysics, UT Southwestern Medical Center, Dallas, TX 75390, USA; Lyda Hill Department of Bioinformatics, UT Southwestern Medical Center, Dallas, TX 75390, USA

## Abstract

The actin cortex plays a large role in regulating the dynamic organization of cell surface receptors, which in turn regulates their signaling. However, many receptors have short intracellular domains and no known link to cortical actin. In this work, we identified the β_1_-integrin subunit and several tetraspanins – CD9, CD81 and CD151 – as part of the hitherto unknown molecular link between the surface receptor CD36 and cortical actin. We found that CD36 in vascular endothelial cells is recruited into complexes/nanodomains containing these proteins, with stronger recruitment near the cell edge. Perturbing this recruitment via the mutation G12V in the N-terminal transmembrane domain of CD36 alters the dynamic organization of CD36 on the vascular endothelial cell surface and weakens its coupling to cortical actin dynamics. Moreover, perturbing this recruitment abolishes thrombospondin-1-induced CD36 signaling through the Src family kinase Fyn. Given their many interactions with other transmembrane proteins, tetraspanins and integrins may provide a ubiquitous mechanism for plasma membrane-cortical actin coupling.

## Introduction

CD36 is an integral membrane protein expressed on the surface of many cell types, including microvascular endothelial cells (MVECs), macrophages, microglia, platelets and various epithelia (Silverstein and Febbraio, 2009). It binds diverse ligands, most notably thrombospondin-1 (TSP-1), oxidized low-density lipoprotein, fibrillar β-amyloid and malaria-infected erythrocytes, and functions as a fatty acid transporter (Pepino et al., 2014; Silverstein and Febbraio, 2009). It is implicated in various physiological and pathophysiological processes, such as angiogenesis, atherosclerosis, Alzheimer’s disease, immunity, diabetes and obesity (Silverstein and Febbraio, 2009).

Like many receptors, especially (but not limited to) those involved in immunity and inflammation (Chen et al., 2021; Garcia-Parajo et al., 2014; Garcia-Parajo and Mayor, 2024; Goyette et al., 2019; O’Shea and Murray, 2008; Treanor et al., 2010; Wilson et al., 2011), CD36 signaling requires clustering and the formation of signaling complexes (Bamberger et al., 2003; Chen et al., 2022; Githaka et al., 2016; Heit et al., 2013; Jaqaman et al., 2011; Jimenez et al., 2000; McGilvray et al., 2000; Wilkinson et al., 2006; Wong et al., 2016). With its two short intracellular domains – 7 and 13 amino acids at the N and C termini, respectively – which lack signaling motifs and scaffolding domains, clustering and complex formation is critical to bring CD36 together with its downstream signaling partners (Chen et al., 2022; Heit et al., 2013). In MVECs, for example, CD36 clusters are enriched with the downstream effector Fyn (a Src family kinase; SFK), thus enabling efficient signaling upon ligand binding (Githaka et al., 2016).

The dynamic organization of CD36 at the plasma membrane (PM), like that of many receptors (Garcia-Parajo et al., 2014; Garcia-Parajo and Mayor, 2024; Jaqaman and Grinstein, 2012), both in the absence and presence of ligand, depends on actin: perturbing actin reduces the dynamic interactions and clustering of CD36 at the PM, and mutes signaling in response to ligand (Githaka et al., 2016; Jaqaman et al., 2011). However, the short intracellular domains of CD36 are unlikely to interact with cortical actin (CA) directly. Rather, CA most likely exerts its influence on CD36 indirectly through other molecules that interact with CA and with CD36 (as observed for other molecules (Freeman et al., 2018; Goswami et al., 2008; Kusumi et al., 2012)). The clusters/nanodomains/complexes with which CD36 associates are prime candidates for mediating this influence. In this, CD36 probably represents many PM components that are influenced by CA, but primarily indirectly through the influence and action of other PM components.

The β_1_-integrin subunit and the tetraspanins CD9, CD81 and CD151 are strong candidates for mediating the molecular link between CD36 at the PM and CA. They physically interact with CD36, including in MVECs (Heit et al., 2013; Huang et al., 2011; Kazerounian et al., 2011; Miao et al., 2001; Primo et al., 2005; Thorne et al., 2000). They also interact with each other (Berditchevski, 2001; Charrin et al., 2009; Yauch et al., 2000; Zhang et al., 2009) and with the actin cytoskeleton (Bailey et al., 2011; Coffey et al., 2009; Geiger et al., 2001; Kim et al., 2011; Sala-Valdés et al., 2006; Schmidt et al., 2024; Shigeta et al., 2003; Takeda et al., 2007; Zhang et al., 2009). As integrins and tetraspanins interact with many proteins at the PM (Charrin et al., 2014; Charrin et al., 2009; Ivaska and Heino, 2011; Vicente-Manzanares and Sánchez-Madrid, 2018), they likely play a ubiquitous role in PM-CA coupling.

In this work, we tested the hypothesis that CD36-β_1_-Integrin-tetraspanin interactions mediate the link between PM CD36 and CA. We combined advanced cellular imaging and quantitative image analysis with point mutation of CD36 to (i) identify where and to what extent CD36 interacts with β_1_-integrin and the tetraspanins CD9, CD81 and CD151, (ii) inhibit CD36 interactions with β_1_-Integrin and tetraspanins, and (iii) determine the consequences of inhibiting these interactions for CD36-CA coupling, the dynamic organization of CD36, and CD36 signaling.

## Results

### CD36 in MVECs colocalizes with β1-integrin and the tetraspanins CD9, CD81 and CD151, especially near the cell edge

To study the interactions of CD36 with integrins and tetraspanins in their cellular context, we took an imaging and quantitative colocalization approach. We transiently expressed Halo-CD36 in telomerase-immortalized microvascular endothelial (TIME) cells (which expressed little CD36 themselves (Dasgupta et al., 2023)), labeled with JF549-Halo-ligand, and then imaged CD36 together with combinations of immunolabeled β_1_-integrin and tetraspanins CD9, CD81 and CD151 using total internal reflection fluorescence microscopy (TIRFM) (Fig. 1A-E). The imaged molecules were then detected with sub-pixel localization (Aguet et al., 2013; Jaqaman et al., 2008), and, after correcting for registration shift between the different channels, their colocalization relationships were assessed using conditional colocalization analysis (Vega-Lugo et al., 2022). Conditional colocalization analysis allowed us to quantify not only the extent of colocalization between a given pair of molecules, but also whether their colocalization was enhanced by either’s colocalization with a third molecule, providing evidence for cooperativity and complex formation.

**Figure 1.**
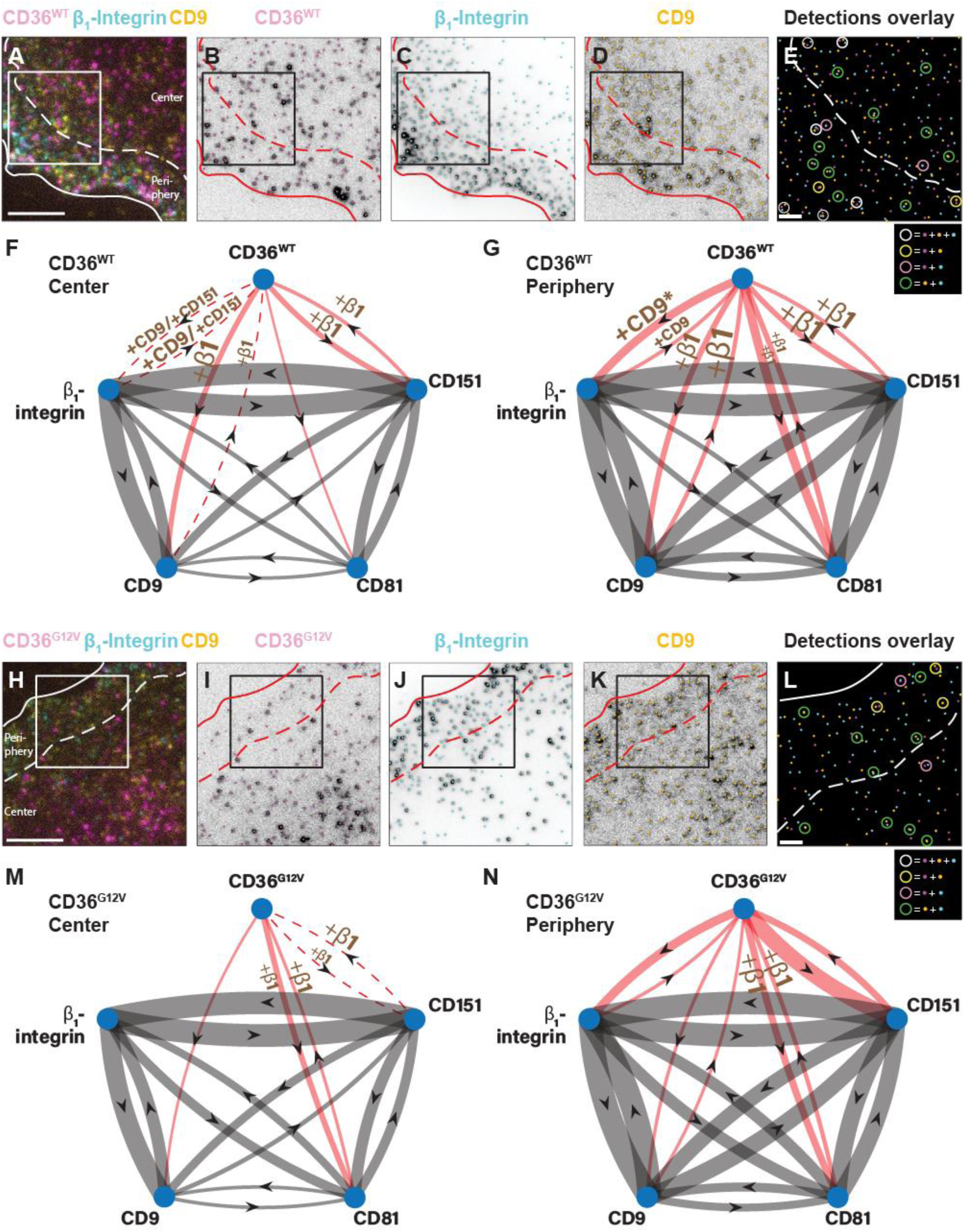
G12V mutation weakens CD36 colocalization with β_1_-integrin and CD9 and alters CD36 colocalization with CD81 and CD151. (**A**) Representative three-color fixed-cell image of CD36 (magenta), β_1_-integrin (cyan) and CD9 (yellow) on the surface of a TIME cell expressing Halo-tagged CD36^WT^ and imaged via TIRFM. Scale bar, 5 μm. (**B-D**) Particle detections (shown as dots) overlaid on the individual channels. (**E**) Overlay of the three-channel detections for the boxed area in A-D, with color coding following that in A-D. Circles highlight colocalization events between the different molecules, following color-coding in the legend at bottom of panel. Scale bar, 1 μm. In A-E, solid lines show segmented cell edge, dashed lines specify boundary between periphery and center. (**F, G**) Representation of the colocalization networks of CD36^WT^, β_1_-integrin, CD9, CD81, and CD151 in the center (**F**) and periphery (**G**) regions. Network arrows involving/not involving CD36 are shown in red/gray, for visual clarity. Arrows point from target molecule to reference molecule and represent colocalization of target with reference. Solid arrows indicate significant colocalization of target with reference (p ≤ 0.05), with arrow thickness proportional to level of significance. “+ molecule name” next to arrow represents condition molecule, and indicates that colocalization of target with reference is significantly enhanced for subset of CD36 colocalized with condition. An exception is CD36 colocalization with β_1_-integrin in the edge region (indicated by * in G), which is significantly enhanced for the subset of CD36 colocalized with CD9 and also for the subset of β_1_-integrin colocalized with CD9. Font size of condition name reflects level of significance of enhancement. Dashed arrows indicate that target colocalization with reference is only significant for the subset of CD36 colocalized with the condition. In (F), +CD9/+CD151 on arrows between CD36 and β_1_-integrin indicates that colocalization is significant for the subset of CD36 colocalized with either CD9 or CD151. Molecules with no connecting arrow do not exhibit significant colocalization. (**H-N**) Same as A-F but for CD36^G12V^. See Table S1 for the significance p-values from which these graphs were generated. See Table 1 for number of cells, number of experimental repeats, and number of objects per channel per ROI used for analysis.

**Table 1.** Dataset size, including average number of objects in each channel per ROI.

| Dataset<br>(Channels<br>1/2/3, i.e.,<br>561/491/640) | No.<br>of<br>cells | No. of<br>repeats | Channel 1 objects per<br>ROI (average) |  |  | Channel 2 objects per<br>ROI (average) |  |  | Channel 3 objects per<br>ROI (average) |  |  |
| --- | --- | --- | --- | --- | --- | --- | --- | --- | --- | --- | --- |
|  |  |  | Whole | Center | Peri-<br>phery | Whole | Center | Peri-<br>phery | Whole | Center | Peri-<br>phery |
| CD36<sup>WT</sup>/<br>$\beta_1$-integrin/<br>CD9 | 32 | 4 | 452 | 317 | 135 | 494 | 296 | 199 | 445 | 304 | 142 |
| CD36<sup>G12V</sup>/<br>$\beta_1$-integrin/<br>CD9 | 36 | 4 | 385 | 293 | 93 | 567 | 350 | 217 | 378 | 257 | 121 |
| CD36<sup>WT</sup>/<br>$\beta_1$-integrin/<br>CD81 | 48 | 6 | 449 | 321 | 136 | 491 | 304 | 184 | 421 | 293 | 128 |
| CD36<sup>G12V</sup>/<br>$\beta_1$-integrin/<br>CD81 | 48 | 6 | 377 | 272 | 105 | 523 | 334 | 189 | 377 | 253 | 124 |
| CD36<sup>WT</sup>/<br>$\beta_1$-integrin/<br>CD151 | 32 | 4 | 363 | 232 | 130 | 606 | 363 | 243 | 328 | 193 | 136 |
| CD36<sup>G12V</sup>/<br>$\beta_1$-integrin/<br>CD151 | 34 | 4 | 315 | 219 | 96 | 598 | 342 | 256 | 362 | 209 | 153 |
| CD9/<br>CD81/<br>CD151 | 32 | 4 | 396 | 311 | 85 | 434 | 317 | 116 | 316 | 229 | 87 |
| CD36<sup>WT</sup>/<br>Caveolin | 36 | 4 | 556 | 424 | 132 | - | - | - | 426 | 406 | 20 |
| CD36<sup>G12V</sup>/<br>Caveolin | 36 | 4 | 574 | 444 | 130 | - | - | - | 424 | 395 | 29 |
| CD36<sup>WT</sup>/<br>pSFK(Y419)/<br>Fyn, - TSP1 | 50 | 6 | 490 | 356 | 134 | 613 | 442 | 173 | 517 | 396 | 121 |
| CD36<sup>WT</sup>/<br>pSFK(Y419)/<br>Fyn, +TSP1 | 50 | 6 | 518 | 375 | 143 | 689 | 506 | 184 | 550 | 426 | 124 |
| CD36<sup>G12V</sup>/<br>pSFK(Y419)/<br>Fyn, - TSP1 | 50 | 6 | 422 | 320 | 103 | 575 | 413 | 162 | 475 | 363 | 113 |
| CD36<sup>G12V</sup>/<br>pSFK(Y419)/<br>Fyn, +TSP1 | 50 | 6 | 458 | 365 | 93 | 627 | 479 | 149 | 517 | 414 | 104 |

We performed conditional colocalization analysis separately for two PM regions, namely periphery (a band of ∼4 μm adjacent to the cell edge), and center (PM areas > ∼4 μm from the cell edge) (Dasgupta et al., 2023). The reasons for this were twofold. First, from a technical standpoint, the imaged molecules were not evenly distributed throughout the imaged PM area; rather, they generally showed higher density at the periphery (Fig. S1). Thus, it was necessary to analyze these two regions separately, to avoid artifactual colocalization due to large-scale molecule co-distribution within the imaged PM area. Second, from a biological standpoint, the periphery had substantively different actin cytoskeleton architecture than the center, with also higher actin density (Fig. S1F) (Ehringer et al., 1999; Mendoza et al., 2015; Schaphorst et al., 1997; Verkhovsky et al., 2003). As our molecules of interest link to actin (directly or indirectly), we analyzed them separately in these two subregions.

As expected from their known physical interactions, we found significant colocalization between CD36, β_1_-integrin and the three tetraspanins, which was stronger at the periphery than in the center (Fig. 1F, G, Fig. S2, Table S1). At the periphery, CD36-tetraspanin colocalization was additionally enhanced for the subset of CD36 colocalized with β_1_-integrin, and CD36-β_1_-integrin colocalization was enhanced for the subset of CD36 colocalized with CD9 (Fig. 1G). In the center, some colocalizations (CD36 with β_1_-integrin, β_1_-integrin with CD36, and CD9 with CD36) were significant only for the subset of CD36 colocalized with a third interaction partner (Fig. 1F). At the same time, β_1_-integrin-tetraspanin colocalization was indifferent to CD36 in both PM regions (no enhancement of pairwise colocalization for the subset colocalized with CD36). These results indicate cooperativity in CD36 colocalization with its interaction partners, suggesting its recruitment to multimolecular complexes or nanodomains containing integrins and tetraspanins (Heit et al., 2013; Kazerounian et al., 2011), which themselves form with or without CD36 (Berditchevski, 2001; Charrin et al., 2009; Yauch et al., 2000; Zhang et al., 2009). Our subregion analysis indicates that CD36 recruitment into these multimolecular complexes/nanodomains is stronger at the periphery than in the center.

### G12V mutation in CD36 reduces its colocalization with β_1_-integrin and CD9 and alters its colocalization with CD81 and CD151

To test whether CD36 recruitment into β_1_-integrin-tetraspanin complexes/nanodomains contributes to the link between CD36 and CA, we sought to disengage CD36 from these complexes/nanodomains. Given integrins’ broad role in actin network dynamics (Bailey et al., 2011; Coffey et al., 2009; Sala-Valdés et al., 2006; Takeda et al., 2007; Toribio and Yáñez-Mó, 2022; Zhang et al., 2009) and the redundancy between tetraspanins (Charrin et al., 2009), we mutated CD36 instead of perturbing integrins or tetraspanins. Recently, the GXXXG sequence motif in the N-terminal transmembrane helix of CD36 (residues 12-16) was shown to mediate CD36-CD9 interactions (Huang et al., 2023). Thus, we generated three CD36 mutants: the double mutant G12V/G16V and two single mutants G12V and G16V. Out of the three, only the G12V mutant expressed on the cell surface (Fig. S3).

Consistent with this motif mediating CD36-CD9 interactions (Huang et al., 2023), three-color imaging and conditional colocalization analysis (as above) revealed that the G12V mutation reduced CD36-CD9 colocalization, both in the center and at the periphery (Fig. 1H-N, Table S1). Interestingly, it also reduced CD36-β_1_-integrin colocalization, even down to insignificance in the center region (Fig. 1H-N, Fig. S2C, D, Table S1) (note that integrins also contain a GXXXG motif, which mediates many PM protein-protein interactions (Schneider and Engelman, 2004; Teese and Langosch, 2015)). The effect of the mutation on CD36-CD81 and CD36-CD151 colocalizations varied by subregion: CD36-CD81 colocalization increased in the center and decreased at the periphery, while CD36-CD151 colocalization showed the opposite trend (Fig. 1M, N, Table S1). The mutation also shifted the cooperativity landscape, abolishing most of the cooperativity in CD36-CD9-β_1_-integrin and CD36-CD151-β_1_-integrin colocalizations, while enhancing it for CD36-CD81-β_1_-integrin colocalization (both center and periphery). All in all, these results provide evidence that the G12V mutation weakens CD36 recruitment into multimolecular complexes/nanodomains of CD9 and β_1_-integrin across the PM. It also alters CD36 interactions with CD151 and CD81, weakening them in some regions (periphery/center for CD81/CD151) while strengthening them in others (center/periphery for CD81/CD151).

### G12V mutation reduces coupling between CD36 mobility type and CA dynamics

The G12V mutation provided us with a molecular handle to test the hypothesis that CD36 recruitment into β_1_-integrin-tetraspanin complexes/nanodomains contributes to the link between CD36 and CA. Previous studies have shown that CD36 mobility (underlying the dynamic organization of CD36) at the PM is influenced by CA (Dasgupta et al., 2023; Jaqaman et al., 2011). Therefore, if CD36-β_1_-integrin-tetraspanin complexes/nanodomains contribute to the link between CD36 and CA, we expect that the mobility of CD36^G12V^ to be less coupled to CA architecture and dynamics than that of CD36^WT^.

To this end, we employed our previously developed live-cell single-molecule imaging–fluorescent speckle microscopy (SMI-FSM) approach (Dasgupta et al., 2023). We expressed Halo-CD36^WT^ or Halo-CD36^G12V^ in TIME cells stably expressing low levels of mNeonGreen-actin (TIME-mNGrActin cell line), such that the CA network appeared as a collection of fluorescent speckles when imaged via TIRFM (Danuser and Waterman-Storer, 2006). We performed live-cell SMI of JF549-labeled CD36 at 10 Hz (WT or mutant), simultaneously with FSM of CA at 0.2 Hz (Dasgupta et al., 2023) (Videos S1-S4). We then (i) processed CD36 SM tracks to obtain tracklets localized in time and space and in terms of their motion type, (ii) processed CA speckles to remove artifactual appearances and movement, and (iii) matched CD36 tracklets to neighboring CA speckles spatially and temporally (Dasgupta et al., 2023; Jaqaman et al., 2008; Ponti et al., 2003). With this, we obtained for every CD36 tracklet (WT or mutant) its mobility properties and the properties of the CA speckles in its vicinity, allowing us to analyze the coupling between the two.

To describe CD36 tracklet mobility, we employed divide-and-conquer moment scaling spectrum (DC-MSS) transient diffusion analysis, where the mobility type of a tracklet is reflected by its MSS slope (Dasgupta et al., 2023; Ferrari et al., 2001; Vega et al., 2018). To assess the extent of coupling between WT or mutant CD36 mobility type and CA architecture and dynamics, we performed multivariate regression (MVRG) analysis of tracklet MSS slope on CA speckle properties (Dasgupta et al., 2023). MVRG analysis describes the (mathematical) dependence of a tracklet property (such as MSS slope) on the CA speckle properties of interest. We did the analysis on a per cell basis, utilizing four subsets of SM tracklets, grouped by their PM location (center or periphery) combined with their local CA speckle density (low or high, i.e. below or above the average speckle density = 1.3 speckles/µm^2^) (Dasgupta et al., 2023).

We performed MVRG analysis of the MSS slope on various CA speckle properties: displacement (reflecting CA network fluctuations), density and intensity (reflecting CA network density) and lifetime (reflecting CA network stability). Only displacement and density yielded significant MVRG coefficients, as in our previous analysis of SM diffusion coefficient (Dasgupta et al., 2023). However, because the MVRG coefficients on density showed the expected positive/negative values at low/high CA speckle densities (Fig. S4), which are most likely due to the discrete nature of speckles and spatial sampling by SM tracklets (Dasgupta et al., 2023), we focused our analysis on the CA speckle displacement component.

CD36^WT^ MSS slope showed a significant negative MVRG coefficient on CA speckle displacement for all subgroups (Fig. 2A, B), consistent with our earlier finding of negative coupling between CA speckle displacement and PM protein mobility (Dasgupta et al., 2023). However, for CD36^G12V^, the MVRG coefficient was insignificant for SMs with low local speckle density, both in the center and at the periphery (Fig. 2A, B). The CD36^G12V^ MVRG coefficient magnitude was also reduced for SMs with high local CA speckle density in the center. The stronger difference between CD36^WT^ and CD36^G12V^ at low local CA speckle density (vs. high speckle density) is reminiscent of our previous finding that differences in coupling to CA between molecules with different actin-binding ability manifest themselves more readily at low local CA speckle density (Dasgupta et al., 2023). Importantly, these results indicate that the G12V mutation weakens the coupling between CD36 mobility type and CA, providing evidence that CD36-β_1_-integrin-tetraspanin complexes/nanodomains contribute to the molecular link between CD36 and CA.

**Figure 2.**
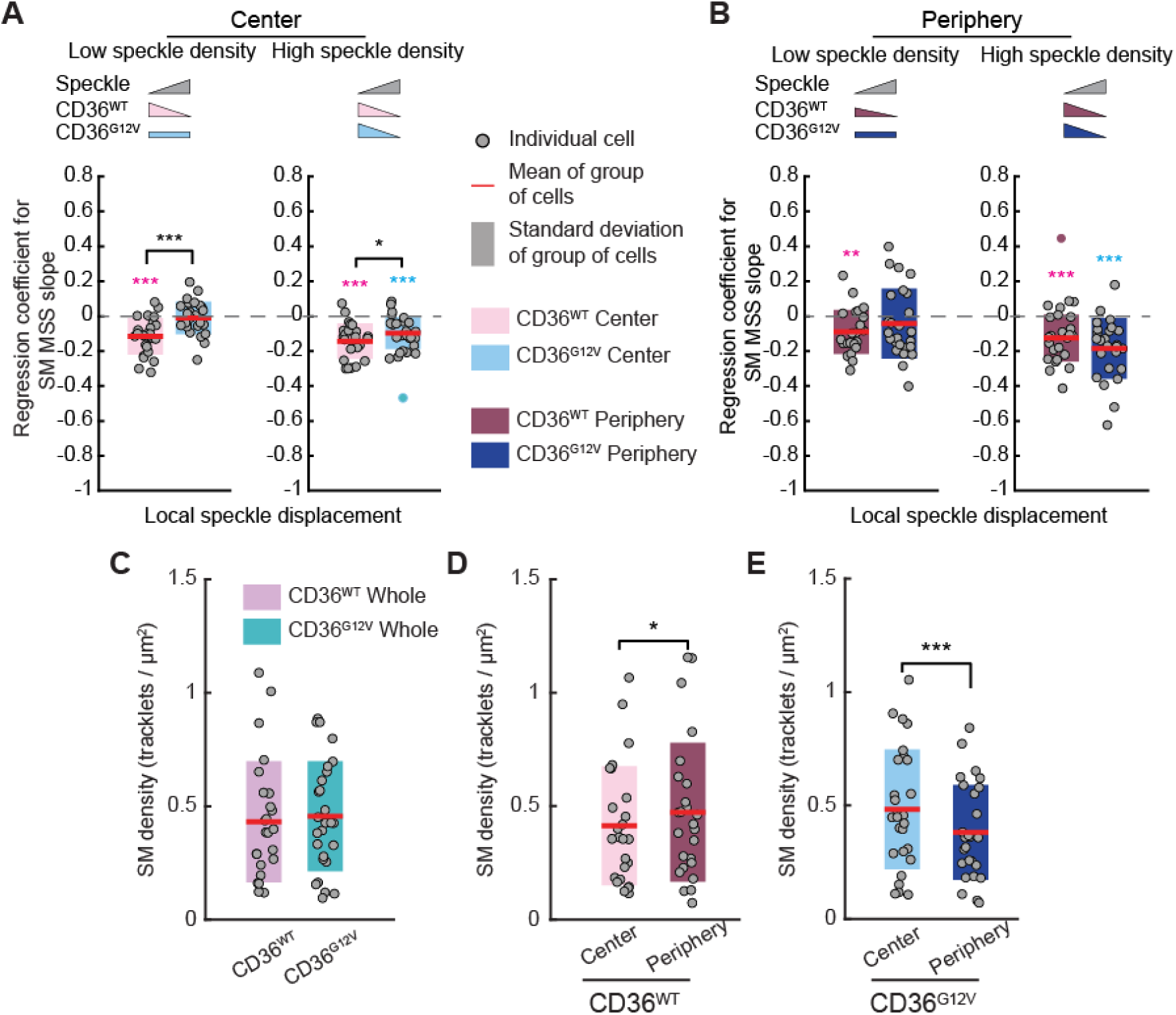
G12V mutation reduces coupling between CD36 mobility type and local CA speckle displacement and redistributes CD36 from the periphery to the center. (**A, B**) Regression coefficients of CD36^WT^ and CD36^G12V^ MSS slope on local CA speckle displacement, from MVRG analysis on speckle displacement and density (see Fig. S4 for corresponding regression coefficients on density). MVRG analysis was performed separately for the four indicated subgroups of SM tracklets based on PM region (center (**A**) and periphery (**B**)) combined with local CA speckle density. Gray circles indicate individual cell measurements; non-gray circles indicate outliers. Red lines and shaded bars show mean and standard deviation, respectively, over group of cells representing WT or mutant. MVRG coefficients significantly different from zero (dashed gray line) are indicated by colored asterisks above the MVRG coefficient plots. Significant differences between WT and mutant are indicated by black asterisks above bracket. For both, *, **, and *** indicate p ≤ 0.05, 0.01, and 0.001, respectively. In all tests, p > 0.05 is not explicitly indicated to reduce clutter. Inset pictograms, visual summary of MVRG results: triangle slope and direction represent significance and sign (positive or negative) of MVRG coefficient; rectangles indicate nonsignificant MVRG coefficients. (**C**) SM tracklet density for WT and mutant CD36 in the whole imaged region. (**D, E**) SM tracklet density in the center and periphery subregions for WT (**D**) and mutant (**E**) CD36. In C-E, circles, red lines, shaded bars and black asterisks (or lack thereof) as in A, B. See Table 2 for number of cells, number of experimental repeats, and number of SM tracklets and speckles per ROI used for analysis.

**Table 2.** SMI-FSM dataset size, including average number of SM and speckle datapoints per ROI.

| Dataset | No. of cells | No of repeats | SM tracklets per ROI (average) |  |  | speckle detection per frame per ROI (average) |  |  |
| --- | --- | --- | --- | --- | --- | --- | --- | --- |
|  |  |  | Whole | Center | Periphery | Whole | Center | Periphery |
| CD36<sup>WT</sup> | 25 | 8 | 2340 | 1629 | 708 | 696 | 462 | 205 |
| CD36<sup>G12V</sup> | 27 | 7 | 2386 | 1890 | 495 | 720 | 489 | 216 |

### CD36^G12V^ localizes less in the actin-rich periphery region of the PM than CD36^WT^

Another evidence for the reduced coupling between CD36 and CA due to the G12V mutation came from the distribution of CD36 molecules across the PM. In both fixed and live cells, CD36^WT^ preferentially localized in the periphery region of the PM (Fig. 2C, D; Fig. S1A), where actin density was higher (Fig. S1F). In contrast, CD36^G12V^ preferentially localized in the center region (Fig. 2C, E; Fig. S1G). Since β_1_-integrin and the investigated tetraspanins also predominantly localize at the periphery (Fig. S1B-E), their interactions with CD36 likely retain CD36^WT^ at the periphery, while the loss/reduction of these interactions leads to a redistribution of CD36^G12V^ toward the center. Importantly, these trends are consistent with the hypothesis that the CD36-CA link is weakened by the G12V point mutation, providing further evidence that CD36 recruitment into β_1_-integrin-tetraspanin complexes/nanodomains contributes to this link.

### CD36^G12V^ exhibits a higher fraction of confined diffusion than CD36^WT^, but in larger confinement areas

What are the consequences of perturbing CD36-β_1_-integrin-tetraspanin complexes/nanodomains and weakening the CD36-CA link for CD36 mobility? Unexpectedly, we found that the mutant exhibits more confined diffusion than WT. First, a higher fraction of tracklets was classified as confined for CD36^G12V^ than for CD36^WT^ (∼24% vs. ∼18% on average), at the expense of free tracklets (Fig. 3A). This was both in the center and at the periphery (Fig. 3B, C). Second, free SM tracklets themselves had an overall lower MSS slope in the mutant vs. WT (Fig. 3D), indicating a greater extent of hidden confined diffusion (or immobility) within the tracklets classified as free in the case of CD36^G12V^. This confined diffusion (or immobility) is “hidden” because it happens transiently, at a faster timescale than our imaging temporal resolution (0.1 s). Such short-lived instances of confinement (or immobility) are undetectable by DC-MSS, but they reduce the overall MSS slope of the classified tracklets (Ritchie et al., 2005).

**Figure 3.**
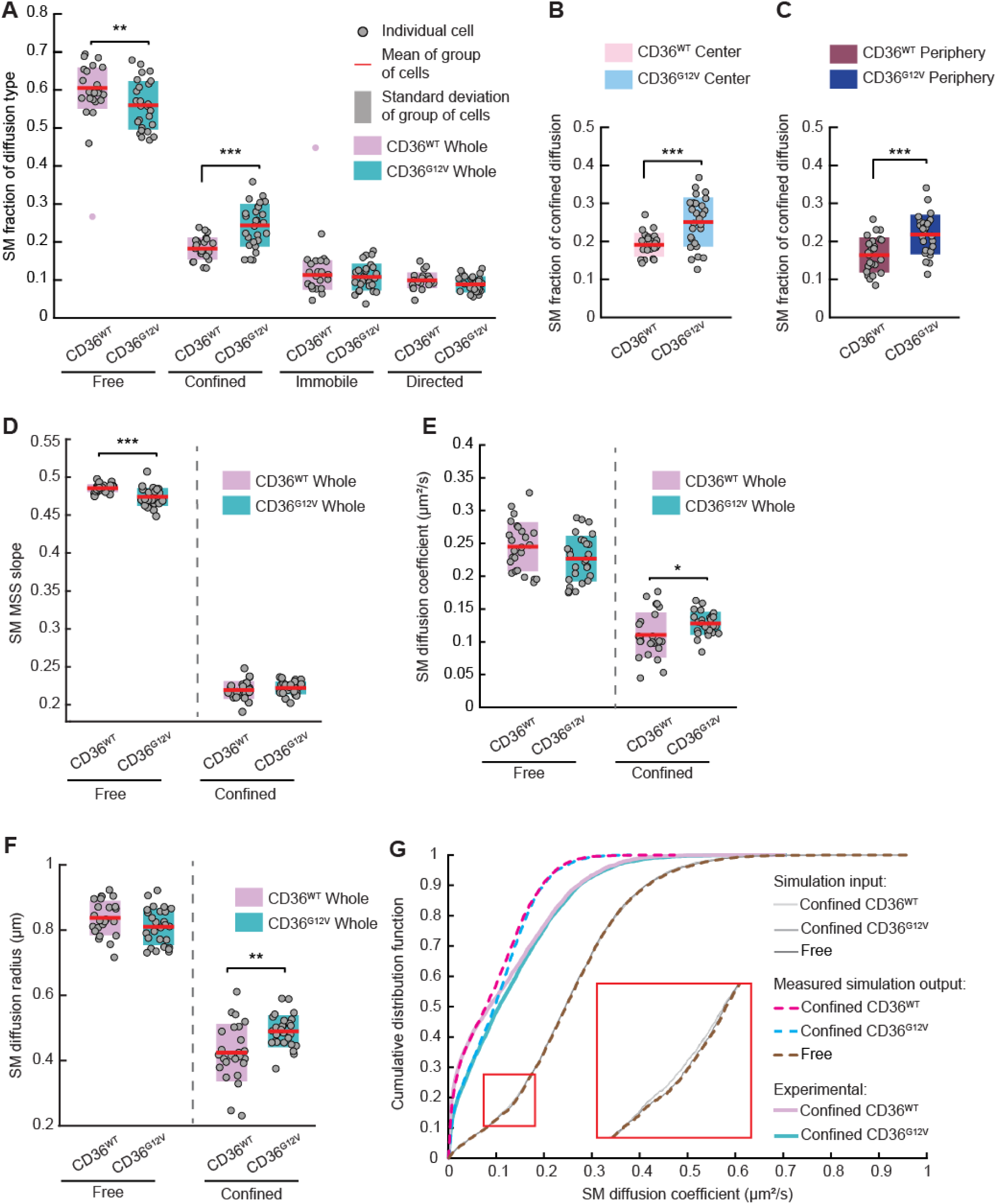
G12V mutation increases fraction of CD36 molecules undergoing confined diffusion, but in larger confinement areas. (**A**) Fraction of CD36^WT^ and CD36^G12V^ tracklets undergoing the indicated motion types in the whole imaged region. (**B-C**) Fraction of CD36^WT^ and CD36^G12V^ tracklets undergoing confined diffusion in the center (**B**) and periphery (**C**) subregions. (**D**) MSS slope for free and confined CD36^WT^ and CD36^G12V^ in the whole imaged region. (**E, F**) Diffusion coefficient (**E**) and diffusion radius (**F**) for free and confined CD36^WT^ and CD36^G12V^ tracklets in the whole imaged region. (**G**) The cumulative distribution function of SM diffusion coefficient as input into simulations (solid gray lines), as measured for simulated free and confined tracks (dashed lines), and as measured experimentally (solid non-gray lines). The red box zooms in on an area for visual aid to distinguish between the highly overlapping gray lines. In all panels but G, circles, red lines, shaded bars, and black asterisks (or lack thereof) as in Fig. 2. See Table 2 for number of cells, number of experimental repeats, and number of SM tracklets and speckles per ROI used for analysis.

Yet the confined CD36^G12V^ tracklets had a larger diffusion coefficient and diffusion radius (reflecting confinement area) than those of CD36^WT^ (Fig. 3E, F). Free SM tracklets showed no difference in their diffusion coefficient or radius between CD36^WT^ and CD36^G12V^ (Fig. 3E, F). The concomitant increase in both the diffusion coefficient and diffusion radius for CD36^G12V^ confined tracklets suggested that these two properties were coupled. This, together with the similar diffusion coefficient of free CD36^WT^ and CD36^G12V^, led us to hypothesize that WT and the mutant have fundamentally the same diffusion coefficient, but that, for the confined subset, different confinement areas between WT and G12V lead to different effective diffusion coefficients at our imaging time resolution (0.1 s).

To test this hypothesis, we simulated two sets of tracks, using for both the diffusion coefficient distribution of free CD36^WT^, combined with the diffusion (i.e. confinement) radius distribution of confined CD36^WT^ for one set and of confined CD36^G12V^ for the other. As a control, we simulated a set of freely diffusing tracks, also using the diffusion coefficient distribution of free CD36^WT^ tracklets. In all simulations, track lifetimes ranged from 20 to 40 time points, sampled at 0.1 s, similar to experimental tracklets. We then applied MSS diffusion analysis to the simulated tracks, similar to experimental tracklets.

Consistent with our hypothesis, the calculated diffusion coefficient of simulated confined CD36^G12V^ tracks was shifted toward higher values compared to that of simulated confined CD36^WT^ tracks (Fig. 3G). For the free diffusion tracks, the calculated diffusion coefficient matched the input diffusion coefficient, as expected (Fig. 3G). Remarkably, not only did the simulated confined diffusion coefficients follow the expected trend between CD36^WT^ and CD36^G12V^, but the diffusion coefficient values matched those measured experimentally for confined tracklets in the lower half of the distribution, which is where the separation between CD36^WT^ and CD36^G12V^ is largest (Fig. 3G). The discrepancy in the upper half of the distribution (where there is little difference between WT and mutant) probably stems from the simplified nature of the simulations, where simulated confined tracks were confined at all times, while experimental confined tracklets probably contained some hidden free diffusion within them, thus increasing their effective diffusion coefficient.

Overall, these results indicate that perturbing CD36-β_1_-integrin-tetraspanin complexes/nanodomains and weakening the link between CD36 and CA lead to mis-regulation of CD36 mobility, such that the balance between free and confined diffusion is shifted toward the confined state. At the same time, the confined state in the mutant spans a larger confinement area, suggesting a weaker “force of confinement.”

### The diffusion properties of confined CD36^G12V^ molecules are less coupled to CA dynamics than those of CD36^WT^

Previously, we found that the diffusion coefficient of confined CD36 tracklets has stronger coupling (larger MVRG coefficient magnitude) to CA speckle displacement than that of free CD36 tracklets (Dasgupta et al., 2023). This implies that confined CD36 tracklets are more strongly linked to CA than free tracklets. Is this still the case for CD36^G12V^? Our observation of reduced coupling between tracklet mobility type and CA for CD36^G12V^ (Fig. 2A, B), and the different confinement area for CD36^G12V^ (Fig. 3E), suggest that the factors underlying CD36^WT^ and CD36^G12V^ confinement are different, with CA playing a smaller role in the mutant case. To test whether confined tracklets were indeed less linked to CA in the G12V case when compared to WT, we performed MVRG analysis of their diffusion properties, namely diffusion coefficient and diffusion radius. We performed the analysis for four subsets of tracklets, grouped by their diffusion type (free or confined) combined with their local CA speckle density (low or high) (Dasgupta et al., 2023).

As expected (Dasgupta et al., 2023), CD36^WT^ diffusion coefficient showed significantly negative MVRG coefficients on CA speckle displacement (Fig. 4A, Fig. S4). The MVRG coefficients for diffusion radius paralleled those for diffusion coefficient (Fig. 4B, Fig. S4). Moreover, for CD36^WT^, the MVRG coefficients of confined tracklets were significantly stronger than those for free tracklets (Fig. 4A, B). Compared to CD36^WT^, the MVRG coefficients for confined CD36^G12V^ tracklets were significantly weaker – at both low and high local speckle density in the case of the diffusion radius (Fig. 4B) and at low local speckle density in the case of the diffusion coefficient (Fig. 4A). In addition, the difference in coupling strength between confined and free tracklets was abolished in the case of the diffusion radius (Fig. 4B) and reduced in the case of the diffusion coefficient at low local CA speckle density (Fig. 4A). The MVRG coefficients for free tracklets were not affected by the mutation, probably because freely diffusing tracklets were less linked to CA in the first place.

**Figure 4.**
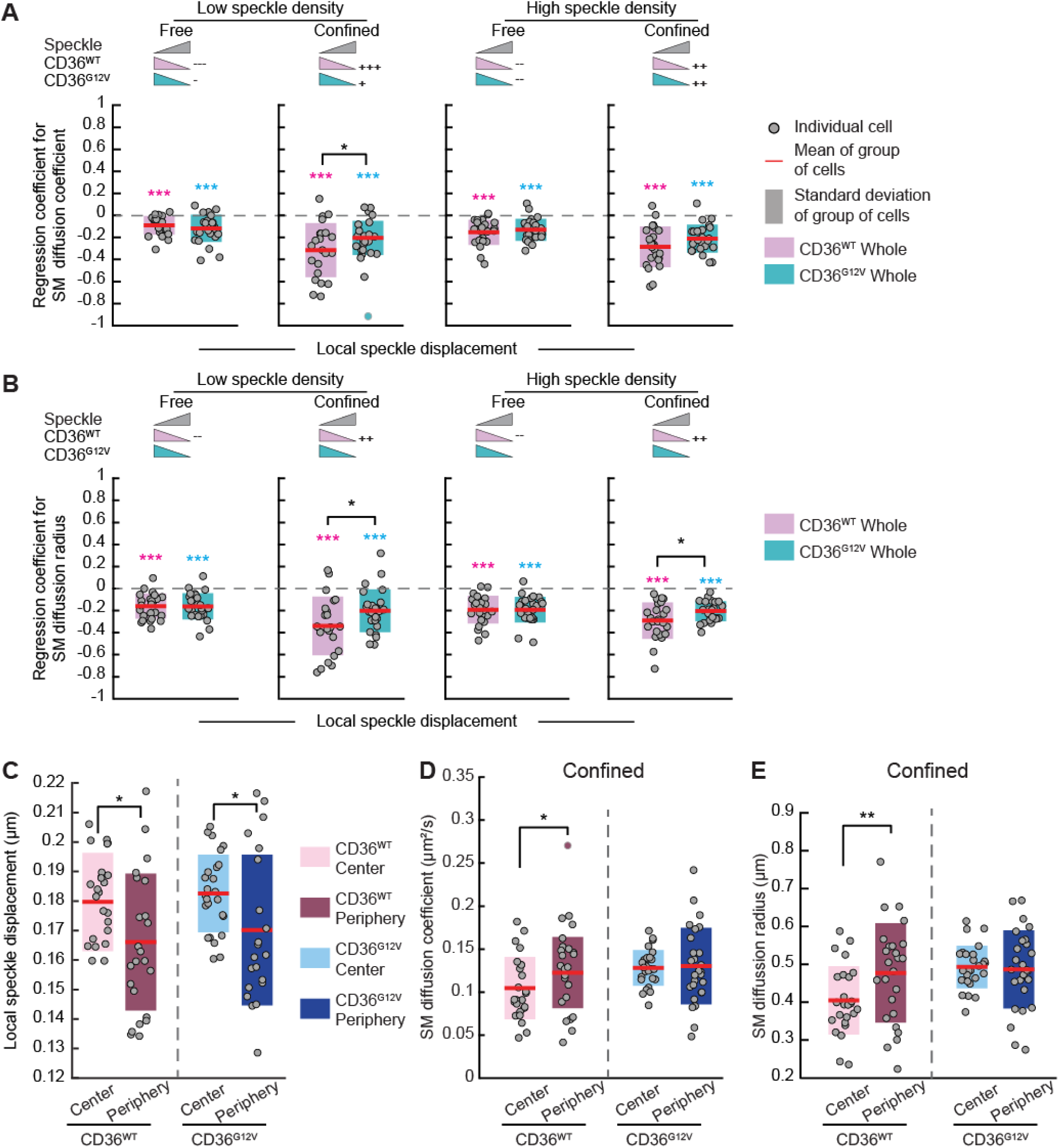
Confined CD36^G12V^ molecules exhibit less coupling to CA speckle displacement than confined CD36^WT^ molecules. (**A, B**) Regression coefficients of CD36^WT^ and CD36^G12V^ diffusion coefficient (**A**) and diffusion radius (**B**) on local CA speckle displacement, from MVRG analysis on speckle displacement and density (see Fig. S4 for corresponding regression coefficients on density). MVRG analysis was performed separately for the four indicated subgroups of SM tracklets based on local speckle density and diffusion type. Next to the pictograms, +, ++, +++/−, --, --- indicate significant differences (greater/smaller; p ≤ 0.05, 0.01, and 0.001, respectively) between confined and free MVRG coefficient magnitudes at the same CA speckle density level for each type of molecule. p > 0.05 is not explicitly indicated to reduce clutter. (**C**) Local CA speckle displacement magnitude at CD36^WT^ and CD36^G12V^ molecules in the center and periphery regions. (**D, E**) Diffusion coefficient (**D**) and diffusion radius (**E**) of confined CD36^WT^ and CD36^G12V^ tracklets in the center and edge regions. Circles, red lines, shaded bars, asterisks (or lack thereof) and pictograms (the latter in A, B) as in Fig 2. See Table 2 for number of cells, number of experimental repeats, and number of SM tracklets and speckles per ROI used for analysis.

In addition to the reduction in MVRG coefficient magnitudes, differences in the diffusion properties of confined tracklets between the center and periphery were abolished in the mutant. Specifically, for CD36^WT^, the diffusion coefficient and diffusion radius of confined tracklets in the center were significantly lower than their counterparts at the periphery, concomitant with the higher local CA speckle displacement in the center compared to the periphery (Fig. 4C-E). These differences between the center and periphery are consistent with the negative relationship between diffusion coefficient/radius and CA speckle displacement (Fig. 4A, B and (Dasgupta et al., 2023)). For CD36^G12V^, however, although it experienced higher local CA speckle displacement in the center compared to the periphery like WT (Fig. 4C), the diffusion coefficient and radius of its confined tracklets were similar between the two regions (Fig. 4D, E).

The reduced coupling between CA dynamics and the diffusion properties of confined tracklets for CD36^G12V^ – manifested in both reduced MVRG coefficients and a lack of difference in diffusion properties between the center and periphery – is consistent with CA playing a smaller role in the confinement of CD36^G12V^ compared to CD36^WT^. This provides further evidence for the weakened link between CD36 and CA upon perturbing CD36 recruitment into β_1_-integrin-tetraspanin complexes/nanodomains, thus implicating them in linking CD36 to CA.

### G12V mutation alters CD36 localization in caveolae and signaling at the PM

As CD36 lacks any known signaling domains or motifs, its organization at the PM is thought to be important to bring it together with signaling partners, whether they be other cell surface receptors or intracellular signaling partners, such as SFKs (Chen et al., 2022; Githaka et al., 2016). CD36 has been shown to signal through various lipid- and protein-based nanodomains and complexes, such as lipid rafts, including caveolae (Febbraio et al., 2001), and multimolecular complexes composed of integrins, tetraspanins, and other proteins (Chu et al., 2013; Heit et al., 2013; Kazerounian et al., 2011). With its altered dynamic organization, we hypothesized that CD36 signaling and function would be compromised (or at minimum altered) due to the G12V mutation.

CD36 localization in caveolae, which are highly abundant in vascular endothelial cells (Frank et al., 2003; Luse et al., 2023), has been shown to regulate various CD36 functions, such as fatty acid uptake, LDL endocytosis and eNOS activation (Gerbod-Giannone et al., 2019; Peche et al., 2023; Uittenbogaard et al., 2000). To investigate whether perturbing the dynamic organization of CD36 via the G12V mutation affected its localization in caveolae, we co-imaged CD36 (WT or mutant) with caveolin-1 as a marker for caveolae in fixed cells (Fig. 5A) and then analyzed CD36 colocalization with the detected caveolin-1 puncta. In the “whole” region analysis, i.e. center and periphery together, we found that CD36^G12V^ colocalization with caveolin-1 was higher than that of CD36^WT^ (Fig. 5B). However, we noticed that, for both WT and mutant, caveolin-1 was primarily localized in the center region of the PM (mean caveolin-1 density of 0.5 and 0.1 detections/μm^2^ in center and at periphery, respectively). Given the redistribution of CD36^G12V^ toward the center (Fig. S1A, top row), we speculated that this played a role in the enhanced CD36 colocalization with caveolin-1 in the mutant. Therefore, we performed colocalization analysis for the center and periphery regions separately. We found that CD36 colocalization with caveolin-1 – for both WT and mutant – was significant only in the center (Fig. 5B). Moreover, the extent of CD36 colocalization with caveolin-1 in the center was only very slightly higher for the mutant than for WT (Fig. 5B, Center). This suggests that the redistribution of CD36 toward the center in the case of the mutant underlies the higher overall colocalization of CD36^G12V^ with caveolin-1 (Fig. 5B, Whole). These changes emphasize the important role that the dynamic organization of CD36 - as mediated by its interactions with β_1_-integrin, tetraspanins and CA – plays in regulating CD36 association with PM nanodomains, such as caveolae, that in turn regulate CD36 function.

**Figure 5.**
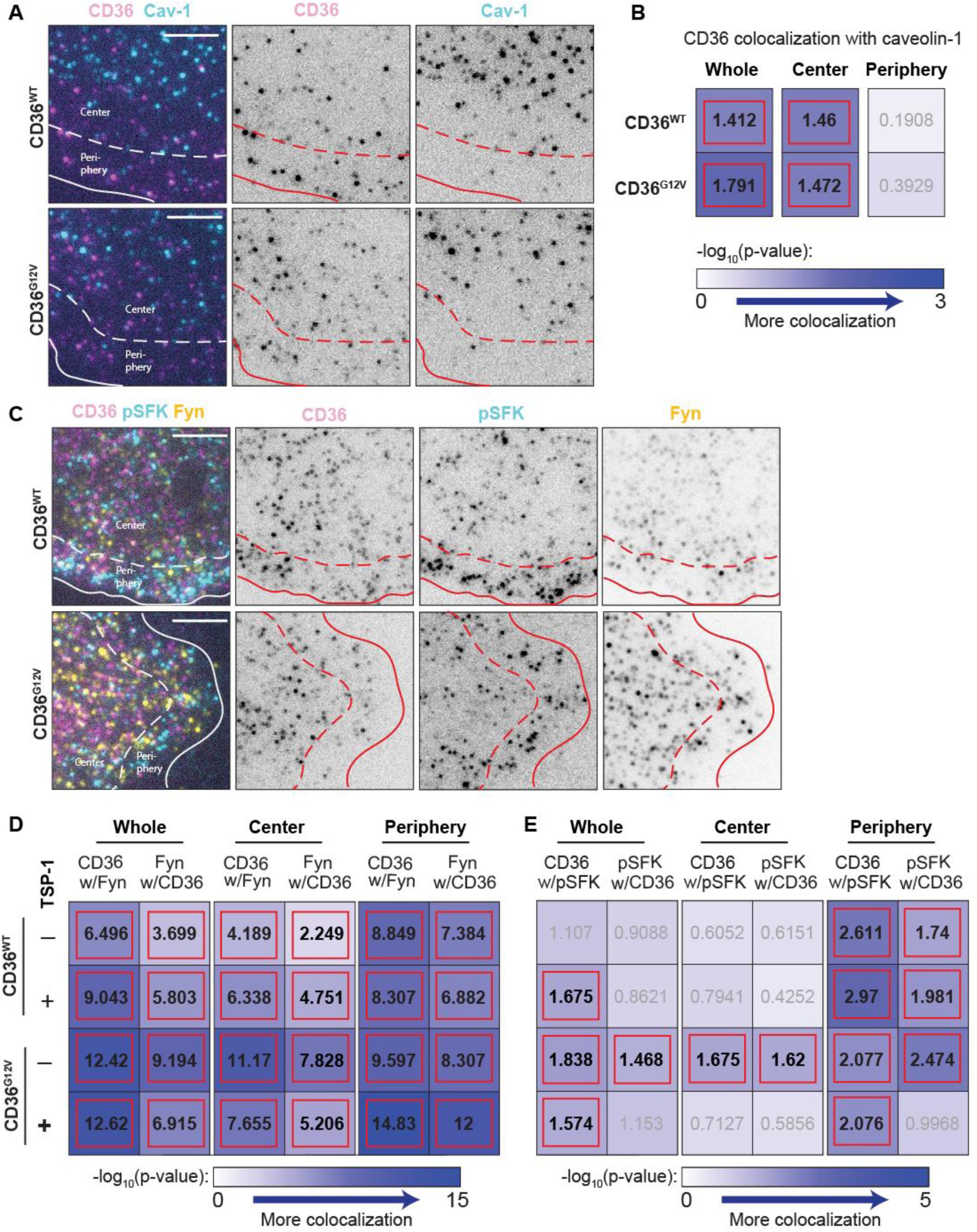
G12V mutation increases CD36 localization at caveolin-1 structures and abolishes CD36 signaling in response to TSP-1. (**A**) Representative two-color fixed-cell images of CD36 (magenta) and caveolin-1 (Cav-1) (cyan) on the surface of a TIME cell imaged via TIRFM, for CD36^WT^ (top) and CD36^G12V^ (bottom). Solid lines show segmented cell edge, dashed lines specify boundary between periphery and center regions. Scale bars, 5 μm. (**B**) −log_10_(p-value) of test for significance of CD36 (WT or mutant) colocalization with Cav-1 puncta in the whole, center and periphery regions. Blue color intensity of each table entry reflects level of significance, as shown in the color bar below table. Values in black and highlighted by solid red outline are significant (p ≤ 0.05; i.e −log10(p) ≥ 1.301); values in gray are not significant. (**C**) Representative three-color fixed-cell images of CD36 (magenta), pSFK(Y419) (cyan), and Fyn (yellow) on the surface of a TIME cell imaged via TIRFM for CD36^WT^ (top) and CD36^G12V^ (bottom). Solid and dashed lines and scale bars as in (A). (**D, E**) −log_10_(p-value) of test for significance of CD36-Fyn colocalization (**D**) and of CD36-pSFK(Y419) colocalization (**E**), for WT and mutant CD36 in the whole, center, and edge regions, in the absence or presence of TSP-1. Details as in (B). See Table 1 for number of cells, number of experimental repeats, and number of objects per channel per ROI used for analysis.

Next, we investigated whether CD36 signaling was perturbed by the G12V mutation. Previous studies have shown that, in MVECs, CD36 binding to its ligand TSP-1 leads to activation of the SFK Fyn (Dawson et al., 1997; Githaka et al., 2016; Jimenez et al., 2000). Therefore, we co-imaged CD36 (WT or mutant) with an antibody against phosphorylated tyrosine 419 in SFKs as well as total Fyn (Fig. 5C), either in unstimulated cells or cells stimulated with TSP-1 (10 nM for 10 min). This imaging-based assay allowed us to measure SFK activation specifically at CD36 molecules (in the form of CD36-pSFK colocalization; (Githaka et al., 2016)) in the context of center vs. periphery PM regions. As expected (Githaka et al., 2016), we found significant CD36^WT^-Fyn colocalization already in the basal state, which was enhanced in the presence of TSP-1 (Fig. 5D, Whole). Center vs. periphery analysis revealed stronger CD36^WT^-Fyn colocalization at the periphery than in the center, irrespective of TSP-1 exposure (Fig. 5D, Center, Periphery). Of note, the colocalization trend was opposite of the trend in Fyn abundance, which was lower at the periphery (0.55 detections/μm^2^ on average) and higher in the center (0.67 detections/μm^2^ on average). Interestingly, Fyn activation at CD36 molecules (CD36-pSFK colocalization) also occurred primarily at the periphery (Fig. 5E, Center, Periphery). The basal level of activation at the periphery was not sufficient to yield significant overall activation, but the increase in activation in the presence of TSP-1 resulted in a significant overall response (Fig. 5E, Whole).

The effect of the mutation on CD36 colocalization with Fyn and on signaling was multifaceted. In contrast to CD36^WT^, CD36^G12V^-Fyn colocalization was stronger in the center than at the periphery in the basal state (Fig. 5D, Center, Periphery). Additionally, there was an elevated level of SFK activation at CD36^G12V^ molecules in the center in the basal state, unlike all other conditions (Fig. 5E, Center). Combined with Fyn activation at CD36^G12V^ molecules at the periphery (Fig. 5E, Periphery; similar to WT), this led to high levels of overall SFK activation at CD36 molecules in the basal state for the G12V mutant (Fig. 5E, Whole). In the background of this high level of SFK activation, TSP-1 addition not only did not lead to further activation, but it rather led to an overall reduction in SFK activation at CD36^G12V^ (Fig. 5E). The center activation was abolished, while the periphery activation was reduced. The extent of CD36-Fyn colocalization went back to being higher at the periphery, similar to WT (Fig. 5D). CD36-Fyn colocalization was in fact the highest for CD36^G12V^ in the presence of TSP-1, yet the level of Fyn activation around CD36 was the lowest, strongly indicating the impairment of CD36^G12V^ signaling in response to TSP-1.

## Discussion

In this work, we have delineated parts of the largely unknown molecular link between the cell surface receptor CD36 and CA, which has been shown to regulate CD36 dynamic organization and signaling. By combining point mutation of CD36 with quantitative analysis of CD36 localization, interactions, mobility and coupling to CA in MVECs, we have identified the β_1_-integrin subunit and multiple tetraspanins as part of this molecular link. Specifically, a mutation in CD36 (G12V) that weakens CD36 association with β_1_-integrin and CD9 and alters CD36 association with CD81 and CD151 (Fig. 1) leads to reduced coupling between the dynamic organization of CD36 at the PM and CA dynamics (Figs. 2, 4). These altered interactions and weakened link to CA are associated with changes in CD36 localization and mobility on the cell surface (Figs. 2, 3, 5), as well as perturbed signaling (Fig. 5).

### Multimolecular complexes including β_1_-integrin and various tetraspanins link CD36 to CA

To the best of our knowledge, this work is the first to provide evidence that β_1_-integrin and the investigated tetraspanins couple the dynamic organization of CD36 at the PM to CA. In previous studies, multimolecular complexes were implicated in CD36 signaling (Heit et al., 2013; Kazerounian et al., 2011); now, they have acquired an additional function. The role of β_1_-integrin and CD9 in mediating the coupling between CD36 and CA is unambiguous, as CD36 colocalization with them is consistently weakened (across the PM) for the G12V mutant. The roles of CD81 and CD151 are less straightforward, because of the PM region (center vs. periphery)-specific effect of the G12V mutation on the extent of CD36 colocalization with them. Nevertheless, given the known network of interactions of these tetraspanins with each other and with β_1_-integrin, as reflected in our study by their strong colocalization with each other (independently of CD36) (Fig. 1), β_1_-integrin and these tetraspanins are expected to mediate the link between CD36 and CA collectively, through complexes and related tetraspanins-enriched nanodomains (Berditchevski, 2001; Charrin et al., 2009; Hemler, 2005; Schmidt et al., 2024).

In terms of CD36 distribution at the PM, we found that CD36 localizes preferentially at the periphery (Fig. S1, Fig. 2), paralleling the localization of its integrin and tetraspanin interaction partners, and the higher density of CA at the periphery (Fig. S1). The higher density of CA at the periphery might itself underlie the preferential localization of β_1_-integrin and the tetraspanins at the periphery, as all of these molecules can physically link to actin (Brakebusch and Fassler, 2003; Sala-Valdés et al., 2006).

It is noteworthy that CD36 colocalization with all its investigated interaction partners is stronger at the periphery than in the center (Fig. 1). This suggests that additional factors – whether they be additional interaction partners, the lipid nano-environment around CD36 complexes/nanodomains, or the CA network itself – enhance and/or stabilize the interactions of CD36 with β_1_-integrin and tetraspanins, and that (some of) these factors are more at play at the periphery than in the center (Berditchevski et al., 2002; Charrin et al., 2003; Schmidt et al., 2024). Interestingly, the extent of colocalization between the different tetraspanins themselves, as well as their colocalization with β_1_-integrin, is also overall stronger at the periphery than in the center. Thus, it is possible that because tetraspanins and integrins form stronger complexes/nanodomains at the periphery (for the reasons mentioned above), CD36 is recruited more successfully into them. Alternatively, factors that disrupt the interactions of CD36 with β_1_-integrin and tetraspanins may be more at play in the center than at the periphery, thus reducing colocalization in the center.

A case at hand, our studies indicate anti-correlation between the extent of CD36 interactions with β_1_-integrin and tetraspanins (more at the periphery, less in the center) and the extent of CD36 localization in caveolae (only in the center, none at the periphery) (Figs. 1, 5). This is consistent with previous work showing that CD36 forms at least two multimolecular complexes, consisting of different proteins and lipids (Kazerounian et al., 2011). Of note, caveolae also link to CA (Echarri and Del Pozo, 2015), thus providing another mechanism linking CD36 to CA. But, having a different molecular nature, the strength and nature of this link is expected to be different from that provided by multimolecular complexes containing β_1_-integrin and tetraspanins.

In terms of CD36 mobility, our results suggest a multifaceted role for CA in regulating CD36 dynamics, probably because of the links between CA and other factors that themselves regulate CD36 organization (such as caveolae, as discussed above). As in our previous work (Dasgupta et al., 2023), our analyses here indicate that, under WT conditions, confined CD36 molecules exhibit stronger coupling to CA than freely diffusing molecules (Fig. 4). This implicates CA in confining (directly or indirectly) CD36 movement at the PM. However, when the link between CA and CD36 is (partly) cut due to the G12V mutation – as evidenced by reduced MVRG coefficients (some becoming insignificant) for CD36^G12V^ MSS slope, diffusion coefficient and diffusion radius on CA speckle displacement (Fig. 2, Fig. 4) – CD36 mobility at the PM does not increase. Rather, a larger fraction of CD36 tracklets exhibits confined diffusion (Fig. 3). At the same time, the strength of confinement is less (larger diffusion radius and consequently diffusion coefficient; Fig. 3), and the coupling of the confined tracklets to CA is lower, becoming similar to the coupling of free tracklets (Fig. 4). This suggests that the confined tracklets in the CD36^G12V^ mutant are (at least partly) different from those in the WT case, and are due to other factors that influence CD36 at the PM. As above, these factors could be lipid-based or other PM nanodomains, or other interaction partners, many of which interact with CA.

### CD36 organization, as mediated by multimolecular complexes and actin, is important for CD36 signaling

A particular strength of imaging-based studies is that they reveal subcellular spatial information that is difficult to obtain otherwise (if at all possible). Here, we found that at least the initial steps of CD36 signaling upon binding its ligand TSP-1 are spatially compartmentalized, occurring primarily in the periphery region of the PM (Fig. 5). We found that CD36-Fyn colocalization is much stronger at the periphery than in the center, and CD36-pSFK colocalization, reflecting Fyn activation at CD36 molecules, occurs primarily at the periphery. In the absence of ligand, there is a basal level of CD36-pSFK colocalization at the periphery, suggesting weak tonic signaling (Githaka et al., 2016; Treanor et al., 2010). Upon TSP-1 addition, CD36-pSFK colocalization increases, to the extent that there is significant CD36-pSFK colocalization even when analyzing the imaged PM area as a whole (which is not the case in the basal state), indicating Fyn activation in response to CD36-TSP-1 binding. Localized Fyn activation might increase the efficiency of CD36 signaling that regulates cell migration (Chu et al., 2013; Dawson et al., 1997).

There are multiple, potentially coexisting, mechanisms by which Fyn activation could occur primarily at the periphery. The stronger complexes at the periphery, including stronger Fyn colocalization with CD36 (Fig. 5), could provide proper nanoscale organization and/or allow Fyn enough residence time in these complexes/nanodomains to get activated upon CD36 binding to TSP-1 (Githaka et al., 2016; Hsieh et al., 2010; Mugler et al., 2012). Another possibility is that additional interaction partners, present primarily at the periphery, and/or the lipid nano-environment at the periphery are necessary for Fyn activation. Furthermore, CA – which is denser at the periphery and has a different network architecture – could play a role in Fyn activation (Sandilands et al., 2007). Distinguishing between these mechanisms will most likely go hand in hand with determining the hitherto unknown molecular mechanisms that lead to Fyn activation upon CD36 binding to TSP-1.

The G12V mutant helps shed light on some of these potential mechanisms. In the mutant, both in the center and at the periphery, there is a decrease rather than an increase in CD36-pSFK colocalization upon TSP-1 addition, despite elevated levels of CD36-Fyn colocalization (Fig. 5). This speaks for a mechanism involving multiple molecular players to activate Fyn upon TSP-1 binding to CD36; Fyn colocalization with CD36 is not enough – other molecules, and potentially their nanoscale organization, partly mediated by CA, are needed (Githaka et al., 2016). Of note, the more striking difference in terms of CD36 colocalization with its interaction partners at the periphery between mutant and WT is not the reduction in colocalization per se, but the loss of colocalization cooperativity in the mutant (Fig. 1), especially for β_1_-integrin and CD9. This points out the need for multimolecular complexes/nanodomains for Fyn activation.

Curiously, the mutant shows a high level of basal-state CD36-pSFK colocalization in the center, which then drops upon ligand addition (Fig. 5). CD36^G12V^ in the basal state is also the only condition where CD36-Fyn colocalization is stronger in the center than at the periphery. It is possible that the increased colocalization of CD36^G12V^ with CD81 in the center (Fig. 1M) allows CD36^G12V^ to engage with new molecules that bring it together with Fyn and mediate Fyn activation independently of TSP-1. It is also possible that the heavily reduced/abolished colocalization of CD36^G12V^ with β_1_-integrin, CD9 and/or CD151 in the center leads to a loss of down-regulators of SFKs in its vicinity, such as CSK (which has been shown to interact with integrins (Maldonado and Leyton, 2023; Obergfell et al., 2002)), resulting in increased Fyn activation. It is tempting to speculate that addition of TSP-1, which binds both CD36 and β_1_-integrin (Chen et al., 2000), brings CD36^G12V^ and β_1_-integrin together, thus restoring some aspects of the WT nanoenvironment around CD36^G12V^, and consequently reducing Fyn over-activation at CD36^G12V^ molecules. Yet just bringing CD36 and β_1_-integrin together by TSP-1 seems not enough, as Fyn activation at CD36 at the periphery is lower for the mutant than for WT, again highlighting the cooperative and multimolecular nature of Fyn activation by CD36.

In conclusion, our work identifies molecules (the β_1_-integrin subunit and the tetraspanins CD9, CD81 and CD151) and PM nanodomains (tetraspanin-enriched nanodomains and caveolae) that contribute to the molecular link between CD36 on the cell surface and CA. Perturbing the interactions between CD36 and its molecular partners is associated with altered CD36 dynamic organization and signaling. Our study highlights the multifaceted nature of PM-CA crosstalk, due to the involvement of multiple interacting and interdependent players. The players that we have identified – integrins, tetraspanins, caveolae – interact with many other PM components. Therefore, the principles learned from our study about the mechanisms linking CD36 to CA and their role in regulating CD36 dynamic organization and signaling are most likely translatable to many other cell surface receptors.

## Materials and Methods

### Cell lines and cell culture

Human telomerase-immortalized microvascular endothelial cells (TIME cells, ATCC, Manassas, VA), and TIME cells stably expressing low levels of mNeonGreen-Actin (TIME-mNGrActin; (Dasgupta et al., 2023)), were used for this study. TIME cells at passage 15-55 and TIME-mNGrActin cells at passage 15-24 were grown in ATCC’s vascular cell basal medium supplemented with microvascular endothelial cell (MVEC) growth kit-VEGF (Catalog No. 50-238-2276), 12.5 µg/mL blasticidine (Thermo Fisher Scientific, Catalog No. A11139-03) and 0.1X of antibiotic-antimycotic (Fisher Scientific, Catalog No. MT30004CI) for 48 h at 37°C + 5% CO_2_ until reaching 80-90% confluency. At these passages, TIME cells express very little endogenous CD36, thus limiting CD36 expression to that from the transfected plasmid (Dasgupta et al., 2023). Whether for fixed-cell imaging (TIME cells) or live-cell SMI-FSM (TIME-mNGrActin cells), 7.4 x 10^4^ cells were seeded on fibronectin-coated (10 µg/mL, Millipore Sigma, Burlington, MA), base/acid cleaned, 0.17 mm (no. 1.5) glass bottom dishes with 14 mm glass diameter (MatTek, Ashland, MA).

### Plasmids

Wildtype (WT) CD36 was fused to HaloTag at its N-terminus (CD36^WT^) (Dasgupta et al., 2023). Starting with this plasmid, single point mutations of Glycine (G) amino acid at the 12^th^ or 16^th^ position to Valine (V) (CD36^G12V^, CD36^G16V^) and double point mutation of the two positions (CD36^G12V/G16V^) were generated by site-directed mutagenesis. For this purpose, the CD36^WT^ plasmid was PCR-amplified with mutation-specific primers (Integrated DNA Technologies; primer details in Table below) and cloned by In-Fusion seamless cloning (Takara, Catalog No. 639648). The mutations were confirmed by sequencing (Fig. S3A) (Plasmidsaurus, Oxford Nanopore Technology).

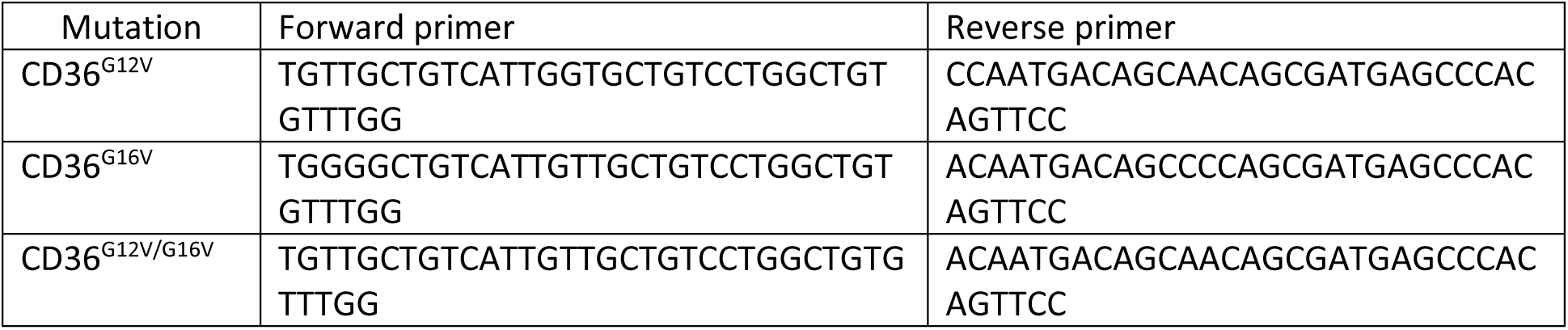

### Transient transfection of CD36

After 18 h of plating on fibronectin-coated glass bottom dishes, TIME cells (for fixed-cell imaging) or TIME-mNGrActin cells (for live-cell SMI-FSM) were transfected with 0.25 μg (/dish) of WT or mutant HaloTag-fused CD36 plasmid (CD36^WT^, CD36^G12V^, CD36^G16V^ or CD36^G12V/G16V^) using Transfex transfection reagent (ATCC, ACS-4005). The medium containing transfection reagent was then replaced with complete culture medium 20 h post transfection. Imaging was performed 2 days post transfection.

### Sample preparation and labelling for fixed-cell imaging

For experiments involving CD36 (except for CD36/pSFK(Y419)/Fyn ± TSP-1; see below for details of these experiments), TIME cells transfected with HaloTag-fused CD36 plasmid (WT or mutant) as described above were incubated for 15 min in complete culture medium containing 15 nM JF549-Halo ligand in the dark. For the CD36 surface expression experiments (Fig. S3), 6 nM JF650-Halo ligand was also added at the same time (although not used for subsequent analysis). The cells were then given a quick wash with sterile DPBS, followed by 15 min incubation with dye-free complete culture medium. All incubations were performed at 37°C + 5% CO_2_. The cells were then washed three times (quick washes) with sterile DPBS. Then, the cells were fixed with 4% paraformaldehyde (PFA) solution made in PBS (Electron Microscopy Sciences, Hatfield, PA, 15710) for 15 min at room temperature (RT), followed by three washes (5 min each) with wash buffer (HBSS + 0.1% normal goat serum (NGS)). For experiments not involving CD36 (3-color imaging of CD9/CD81/CD151), TIME cells were plated on fibronectin-coated glass-bottom dishes for 18 h, and then, without CD36 transfection, they were fixed as described above.

For the CD36/pSFK(Y419)/Fyn ± TSP-1 experiment, the cells were serum-starved for 3 h. After about 2 h of starvation, CD36 (WT or mutant) was labelled with 15 nM JF549-Halo ligand as described above, but in serum free medium. Then the samples were exposed to either 10 nM TSP-1 in HBSS (+ TSP-1) or HBSS alone (-TSP-1, control) for 10 min at 37°C. The cells were then fixed with 4% PFA + 0.1% glutaraldehyde for 20 min at 4°C, followed by quenching (0.1% NaBH_4_ in HBSS) for 7 min at RT, and then three washes (5 min each) with washing buffer (HBSS + 0.1% NGS).

After fixation, there were one or two rounds of primary + secondary (where needed) antibody labeling, depending on the combination of antibodies and their permeabilization requirements (see Table below for details). Each round of labeling was preceded by sample blocking for 15 min or 30 min in blocking buffer (3% BSA and 5% NGS in HBSS). Also, antibodies for labeling, as well as phalloidin, were diluted in blocking buffer. All blocking and labeling were done at RT. The samples were washed three times (5 min each) with wash buffer after each antibody labeling step (whether primary or secondary). Where required, the samples were permeabilized with ice-cold 0.1% Triton X-100 in PBS for 1 min, followed by labelling with antibodies that bind to an intracellular target. Sample permeabilization was always done after labeling all targets accessible without permeabilization. After all rounds of labeling, cells were incubated in imaging buffer (Oxyfluor 1%, Glucose 0.45%, Trolox 2 nM) to reduce photobleaching before and during imaging. Labeling details are in the following table.

|  |  | Round 1 of labeling |  |  | Round 2 of labeling |  |  |
| --- | --- | --- | --- | --- | --- | --- | --- |
|  | Permeabilization | Blocking (min) | Primary antibody | Secondary antibody | Blocking (min) | Primary antibody | Secondary antibody |
| CD36 Halo-JF549 / $\beta_1$-integrin / CD9 | Yes<br>After CD36 and $\beta_1$-integrin<br>Before CD9 | 30 | Anti-$\beta_1$-integrin K20 (Beckman Coulter, IMBULK1B) 1:10,000 1 h | Anti-Mouse-AF488 (Thermo Fisher Scientific, A-11029) 1:1000 15 min | 30 | Anti-CD9-AF647 (sc-13118) 1:25 30 min | - |
| CD36 Halo-JF549 / $\beta_1$-integrin / CD81 | No | 30 | Anti-$\beta_1$-integrin K20 1:100,000 1 h | Anti-Mouse-AF488 1:1000 | 30 | Anti-CD81-AF647 (sc-23962) 1:300 | - |
|  |  |  |  | 15 min |  | 30 min |  |
| CD36 Halo-JF549 / $\beta_1$-integrin / CD151 | No | 30 | Anti-$\beta_1$-integrin K20<br>1:100,000<br>1 h | Anti-Mouse-AF488<br>1:1000<br>15 min | 30 | Anti-CD151-AF647<br>(ab199200)<br>1:250<br>30 min | - |
| CD9 / CD81 / CD151 | Yes<br>After CD81 and CD151<br>Before CD9 | 30 | Anti-CD151-AF647<br>1:250<br>&<br>Anti-CD81-AF488<br>1:100<br>Both for 30 min | - | 30 | Anti-CD9-AF546<br>1:200<br>30 min | - |
| CD36 Halo-JF549 / CD36 Halo-JF650 / Anti-CD36 FA6-152<br>Note: CD36 Halo-JF650 not used for any analysis | No | 15 | Anti-CD36 FA6-152<br>1:400<br>30 min | Anti-Mouse-AF488<br>1:1000<br>15 min | - | - | - |
| CD36 Halo-JF549 / Anti-CD36 FA6-152 / Caveolin-1<br>Note: Anti-CD36 FA6-152 not used for any analysis | Yes<br>After CD36-Halo and anti-CD36 FA6-152<br>Before Caveolin-1 | 15 | Anti-CD36 FA6-152 (ab17044)<br>1:400<br>30 min | Anti-Mouse-AF488<br>1:1000<br>15 min | 30 | Anti-Caveolin-1 (ab32577)<br>1:200<br>30 min | Anti-Rabbit-AF647 (Invitrogen, A32733)<br>1:1000<br>15 min |
| CD36 Halo-JF549 / pSFK(Y419) / Fyn | Yes<br>After CD36<br>Before pSFK(Y419) and Fyn | 15 | Anti-phospho-Src (Tyr419)-AF488 (Thermo Fisher Scientific, 44-660A1)<br>1:500 | Anti-Mouse-AF647 (Thermo Fisher Scientific, A21236)<br>1:1000<br>15 min | - | - | - |
|  |  |  | &<br>Anti-Fyn<br>(ab1881)<br>1:400<br>Both for 1 h |  |  |  |  |
| CD36 Halo-JF549 /<br>Caveolin-1<br>/<br>Phalloidin<br>Note: Only<br>phalloidin<br>channel<br>was used<br>from this<br>dataset | Yes<br>After CD36<br>Before Caveolin-1 and Phalloidin | 30 | Anti-Caveolin-1<br>1:200<br>30 min | Anti-Rabbit-AF488<br>1:1000<br>15 min | Phalloidin-AF647<br>(Thermo Fisher Scientific, A22287)<br>1:20<br>15 min |  |  |

### Fixed-cell imaging

After labeling, the cells were imaged at 37°C using an Olympus IX83 TIRF microscope equipped with a Z-Drift Compensator and a UAPO 100X/1.49 NA oil-immersion TIRF objective (Olympus, Tokyo, Japan). The microscope was equipped with an iXon 888 1k × 1k EMCCD Camera (Andor; Oxford Instruments). With an additional 1.6X magnification in place, the pixel size in the recorded image was 81 nm × 81 nm. Using the Olympus cellSens software, excitation light of 640, 561, and 491 nm from an Olympus CellTIRF-4Line laser system was directed to the sample by a TRF8001-OL3 Quad-band dichroic mirror. Fluorescence of different wavelengths was collected, filtered with emission filters of ET520/40m, ET605/52m, and ET705/72m (Chroma), and projected onto different sections of the camera chip by an OptoSplit III 3-channel image splitter (Cairn Research, Faversham, UK). The different channels were excited and recorded sequentially in the order 640 then 561 then 491, with an exposure time of 99 ms each. Images were acquired with MetaMorph (Molecular Devices, San Jose, CA). Camera EM gain was set to 100 for all acquisitions. The penetration depth was set to 90 nm via the cellSens software (Olympus). Laser powers at sample position (in the widefield illumination configuration) were 2.4, 7.06, and 2 mW for the 491, 561, and 640 nm lasers, respectively.

For every three-channel image, a brightfield snapshot of the imaged cell region was also acquired, to aid with manual delineation (if needed) of the region of interest (ROI) mask for the ensuing analysis.

### Bead preparation and imaging for registration shift correction in fixed cells

To acquire images of Tetraspeck beads (Thermo Fisher Scientific, T7279) for calculating the registration shift between the 3 channels in fixed cell images (491, 561 and 640), Tetraspeck beads were suspended (1:100) in distilled water using a sonicator-water bath for 15 min. The mixed bead sample was then combined with 10 µl poly-L-Lysine (Newcomer supply, Middleton, WI) diluted 1 in 10 µl in nuclease free water, and then plated on a cleaned Mattek dish (as used for cells but without fibronectin coating) for 30 min at RT in the dark The bead sample was imaged using the Olympus IX83 TIRF microscope described above. The penetration depth was set to 90 nm via the cellSens software (Olympus). Laser powers at sample position (in the widefield illumination configuration) were 0.62, 1.76, and 1.43 mW for the 491, 561, and 640 nm lasers, respectively.

### Halo labeling of CD36 for live-cell SMI-FSM

TIME-mNGrActin cells, transfected with WT or mutant CD36 as described above, were incubated for 15 min in complete culture medium with 3 nM JF549-Halo ligand (∼2 days post transfection). Incubation with dye was followed by three quick washes in sterile DPBS. Cells were then incubated for 15 min in dye-free complete culture medium. Incubations were at 37°C + 5% CO_2_. Finally, cells were washed three times (5 min each) with wash buffer (HBSS + 1 mM HEPES and 0.1% NGS). Cells were then incubated in imaging buffer (OxyFluor 1%, glucose 0.45%, Trolox 2 nM) to reduce photobleaching before and during imaging.

### Live-cell SMI-FSM data acquisition

Live TIME-mNGrActin cells expressing CD36^WT^ or CD36^G12V^, both fused to HaloTag and labeled with JF549-Halo ligand as described above, were imaged at 37°C using the Olympus IX83 TIRF microscope described earlier. The videos were acquired with MetaMorph in the stream acquisition mode, with the two channels acquired simultaneously, using the trigger function to control illumination. The 561 channel was used for acquiring SMI videos. These were acquired at 10 Hz for 50 s (501 frames). The 561 channel was triggered to remain open (i.e., continuous illumination) for the entire 501 frames. Simultaneously, the 491 channel was used for acquiring FSM videos. While the 491 channel was acquired also at 10 Hz, the 491 channel was triggered to illuminate the sample only every 50^th^ frame (for 99 ms each). In addition, image acquisition in the 491 channel started 500 frames (50 s) before the start of the 561 channel image acquisition and continued for 500 frames (50 s) after the end of the 561 channel image acquisition (all while illuminating only every 50^th^ frame). Removing the ‘‘empty’’ 491 channel frames (those in the absence of 491 illumination) (using MetaMorph, post video recording), resulted in an FSM time-lapse at 0.2 Hz for 31 FSM frames (150 s), with the middle 11 FSM frames (50 s) corresponding to the 501 SMI frames. The additional FSM frames before and after the simultaneous SMI-FSM period extended the time window of FSM imaging to capture the birth and death of most actin speckles that got matched to SM tracklets in the middle 11 FSM frames, thus allowing accurate speckle lifetime measurement (88% of speckles with known birth and death times). Speckle lifetime reflects the stability of the cortical actin network (longer lifetime = more stable network). Temperature and humidity were maintained during imaging using an environment chamber (Okolab, Otaviano, Italy), maintaining cell viability for the duration of the experiments. The penetration depth was set to 80 nm via the cellSens software (Olympus). Laser powers at sample position (in the widefield illumination configuration) were 3.08 and 7 mW for the 491 and 561 nm laser lines, respectively. Every SMI-FSM movie was preceded by a brightfield snapshot of the imaged cell region to visually check cell viability and to aid with manual delineation (if needed) of the ROI mask for the ensuing analysis.

### Cell segmentation (whole, center, periphery)

In fixed-cell experiments, the cell ROI masks were segmented manually based on the brightfield images and β_1_-integrin channel if available, except for the phalloidin (actin) dataset, where the ROI masks were segmented automatically based on the phalloidin intensity using the u-segment package (https://github.com/DanuserLab/u-segment (Lee et al., 2015; Noh et al., 2022)). In live-cell experiments, the cell ROI masks were generated either within the QFSM package at the thresholding and mask refinement steps (Steps 2 and 3) or they were hand-drawn if thresholding failed (Dasgupta et al., 2023). Starting with these “whole” ROI masks, the “center” ROI masks were created by eroding the whole ROI masks with a disk-shaped structuring element of radius 50 pixels (4.05 µm). The “periphery” ROI masks were then created by subtracting the “center” masks from the “whole” masks. This process resulted in “center” and “periphery” regions being roughly > 4 µm and < 4 µm, respectively, from the segmented cell edge.

### Punctate object detection in fixed cells and Tetraspeck beads

Punctate objects in fixed-cells and Tetraspeck bead experiments were detected using the “point-source detection” particle detection algorithm in u-track (https://github.com/DanuserLab/u-track) (Aguet et al., 2013; Jaqaman et al., 2008). In brief, the algorithm consists of two steps: (i) a convolution/filtering step to determine pixels likely to contain objects and (ii) a Gaussian fitting step to determine the object positions with subpixel localization. With the appropriate, wavelength-dependent standard deviation (1.2, 1.35, and 1.58 pixels for the 491, 561 and 640 channels, respectively), a two-dimensional Gaussian is a good approximation of the microscope’s point spread function (Thomann et al., 2002; Zhang et al., 2007). Largely default parameter values were used, except for in some cases the α-value for determining the significance of detected objects (by comparing the fitted Gaussian amplitude to the local background noise distribution), which was chosen based on visual assessment of the detection results with the goal of minimizing false positives (superfluous detections) and false negatives (missed particles). The default α-value was 0.05, and following are the nondefault cases: β_1_-integrin (491 channel in all experiments), 0.01; CD81 (491 channel in the CD9/CD81/CD151 experiment), 0.01; Caveolin-1 (640 channel), 0.1; and Tetraspeck beads – 561 channel, 10^−6^ & 491 and 640 channels – 0.001. The number of objects detected within the ROI for each fixed-cell dataset is listed in Table 1.

### Registration shift and channel alignment

To increase the accuracy of the ensuing colocalization analysis, the coordinates of detected objects in the three channels were aligned relative to each other. This was achieved by calculating a registration shift between the three channels, using Tetraspeck bead images acquired on the same day or, at most, in the same week. While in the OptoSplit III set up the registration shift between channels was primarily a constant translation across the imaged area, the Tetraspeck beads density was high enough to allow us to calculate the registration shift as a function of (x,y)-coordinates, increasing the accuracy of the registration shift correction. In general, the registration shift between channels was stable and varied very little from week to week. With the registration shift, the alignment between channels was improved from 2-6 pixels (before registration shift) to 0-2 pixels (after registration shift).

In brief, the Tetraspeck bead detections in the three channels were matched between channels using the Linear Assignment Problem (Jaqaman et al., 2008), using a search radius of 9 pixels between 491 and 561 and a search radius of 5 pixels between 640 and 561. The 561 channel was taken as the reference. The matchings from multiple images acquired on the same day (3-5 images) were then utilized together to increase imaged area coverage. Using the matched bead detections, the registration shifts in x and y (Δx and Δy) as a function of position (x and y) were fit using a quadratic polynomial, deemed visually as the most appropriate fit. With this fit, the detections in the fixed-cell multi-channel images were then aligned prior to colocalization analysis.

### Conditional colocalization analysis in different plasma membrane regions of interest (ROIs)

Conditional colocalization analysis of the fixed-cell images was performed as described in (Vega-Lugo et al., 2022). In brief, conditional colocalization analysis quantifies the colocalization of one molecular entity (target) with another molecular entity (reference) and how much this colocalization changes based on either’s colocalization with a third molecular entity (condition). Channel alignment (described above) allowed us to use a relatively small colocalization radius of 2 pixels (= 162 nm) to increase the accuracy of this analysis. Colocalization analysis was done separately for the “whole,” “center,” and “periphery” ROI masks. For each analysis, only the area of the ROI was used for any randomization and other procedures to assess the significance of the observed colocalization measures (Helmuth et al., 2010; Vega-Lugo et al., 2022).

### Statistical testing for colocalization analysis

All statistical testing was as described previously (Vega-Lugo et al., 2022). In brief, for all colocalization analyses, the significance of colocalization is shown in the form of −log_10_(p-value). The p-value for assessing significance was calculated as follows: For p(TwR) (2-way colocalization), the data p(TwR) was compared to its corresponding nullTR (obtained by replacing target objects with points on a grid) using a Wilcoxon rank-sum test. The significance threshold for p(TwR) was set to 0.05. For the conditional colocalization measures (all but p(TwR)), three rank-sum tests were performed to assess significance, namely comparison of each measure to the corresponding nullTR, randC (obtained by randomizing the condition objects, repeated 100 times) and p(TwR). The reported p-value was the maximum (i.e. least significant) of these three tests. For a conditional measure to be significant, its maximum (i.e. reported) p-value had to be < 0.017 (using the Dunn–Sidak correction to obtain a total type-I error of 0.05 for the three tests).

### Live-cell SMI-FSM analysis

SM tracks were constructed from the SMI streams using u-track (version 2.2.1; https://github.com/DanuserLab/u-track)(Jaqaman et al., 2008). The SM detection and tracking parameters were identical to those used previously (Dasgupta et al., 2023). The tracks output by u-track were then processed to remove artifactual merging and splitting events (in addition to those described previously (Dasgupta et al., 2023), very short merge-to-split events lasting <= 2 frames were eliminated as well), and to discard short tracks (duration < 10 time points) and tracks outside the cell mask. SM tracks were then further processed into tracklets localized in space and time and in terms of diffusion type for matching with actin speckles (Dasgupta et al., 2023). SM tracklets’ mobility was analyzed with both divide-and-conquer moment scaling spectrum (DC-MSS) analysis (Vega et al., 2018) and diffusion mode analysis (da Rocha-Azevedo et al., 2020; Jaqaman et al., 2016), although in this work only the DC-MSS results were used. Properties characterizing each SM tracklet were obtained, including (as used in this work):

- Diffusion MSS slope (characterizes diffusion type).
- Diffusion type (free, confined, immobile, or directed).
- Diffusion coefficient.
- Diffusion radius (describes the area covered by the SM tracklet during its lifetime; equivalent to confinement radius for confined SMs).

On the other hand, FSM time-lapse sequences were analyzed using the Quantitative FSM (QFSM) software package (https://github.com/DanuserLab/QFSM) (Danuser and Waterman-Storer, 2006) following the instructions in (Mendoza et al., 2012), and processed to remove “ghost speckles” and artifactual movement, as done previously (Dasgupta et al., 2023). Individual CA speckle properties (e.g., position, displacement, intensity, and lifetime) were then calculated from the speckle tracks output by QFSM software (Dasgupta et al., 2023). After matching actin speckles to SM tracklets temporally and spatially, we calculated the local speckle properties around each SM tracklet by averaging the individual speckle properties (Dasgupta et al., 2023).

For quantifying the mathematical dependence of SM tracklet properties on the matched local speckle properties, linear multivariate regression (MVRG) was performed on an individual cell/ROI (movie) basis. SM tracklet properties were taken as the response (“dependent’’) variables and the associated local speckle properties were taken as the design matrix (‘‘independent variables’’). Property normalization, outlier removal, and MVRG analysis were carried out as described in (Dasgupta et al., 2023). The number of SM tracklets and speckle tracks within the ROI for each live-cell SMI-FSM dataset is listed in Table 2.

### Statistical testing for SMI-FSM

All statistical tests were performed as previously described in (Dasgupta et al., 2023).

In brief, to compare SM or speckle properties, or MVRG coefficients, between CD36^WT^ and CD36^G12V^, a two-sample t-test was performed. To compare SM or speckle properties, or MVRG coefficients, between the Free and Confined subsets of SM tracklets in a cell, or between the Center and Periphery regions of a cell, a paired-sample t-test was performed. For all tests, comparisons with p-values ≤ 0.05, 0.01, or 0.001 were indicated in the figures using *, **, or ***, respectively, while comparisons with a p-value > 0.05 were not marked (to reduce clutter), unless noted otherwise in figure legends.

To determine whether there was a significant relationship between an SM tracklet property and an associated CA speckle property, a one-sample t-test was performed, with the null hypothesis that the single-cell MVRG coefficients followed a normal distribution with mean zero. A relationship was considered significant if the t-test yielded a p-value ≤ 0.05. To assist with visualizing these relationships, the MVRG coefficient plots were annotated with overhead triangle pictograms, where the triangle slopes represent the significance and sign of the MVRG coefficients. Nonsignificant MVRG coefficients were represented with rectangles.

### Simulation of free diffusion and diffusion in confinement areas of different sizes

Three sets of 4,305,000 SM tracks each were simulated. Each set consisted of 4,100 different diffusion parameters (diffusion coefficient and confinement radius, the latter only in the confined diffusion case) × 21 lifetimes (20-40 frames) × 50 repeats. One set (the control) consisted of free diffusion, while two sets consisted of confined diffusion. All simulations utilized the diffusion coefficient distribution of free CD36^WT^ tracklets. The two confined diffusion datasets utilized the confinement radius distribution of confined CD36^WT^ tracklets for one and that of confined CD36^G12V^ tracklets for the other.

2D Brownian motion was simulated as a 2D random walk, with each step in x- and y-direction drawn from a normal distribution with mean 0 and standard deviation equal to √2*D*Δ*t*. D = the diffusion coefficient of a particular simulation. Δt = the simulation timestep = 0.1 s (equivalent to experimental data). For confined diffusion, the particle was simulated to undergo this random walk inside a circular area, with radius R = confinement radius of a particular simulation. The particle was retained inside the circular area by bouncing it off the circle boundary if a step would take it across the boundary.

## Video legends

**Video S1. SMI of HaloTag-labeled CD36^WT^ (left) and CD36^G12V^ (right) labeled with JF549 in TIME-mNGrActin cells.** Video consists of the 49 SMI frames (acquired at 10 Hz) within one FSM interval, and excludes the first and last SMI frames of the interval (which coincide with the FSM frames). Each displayed area is 16.2 × 16.2 μm^2^, cropped from the full acquired image (27.62× 64.8 μm2) for visual clarity.

**Video S2. FSM of CA visualized as mNeonGreen-Actin speckles in the same TIME-mNGrActin cells as in Video S1 expressing CD36^WT^ (left) or CD36^G12V^ (right).** Video consists of 11 FSM frames (acquired at 0.2 Hz) = 50 s (= 10 FSM intervals). Each displayed area is 16.2 × 16.2 μm^2^, cropped from the full acquired image (27.62× 64.8 μm2) for visual clarity.

**Video S3. SM particle tracking.** Zoomed in areas of Video S1 showing particle tracks overlaid on the fluorescence images for CD36^WT^ (left) and CD36^G12V^ (right). Random track colors are used to help distinguish between neighboring tracks. Each displayed area is 8.1 × 8.1 μm^2^ for visual clarity.

**Video S4. FSM speckle detection and tracking.** Zoomed in areas of Video S2 showing speckle detections and tracks overlaid on the fluorescence images for CD36^WT^ (left) and CD36^G12V^ (right). Circles and triangles represent primary and secondary speckle detections (Mendoza, Michelle C et al. “Quantitative fluorescent speckle microscopy (QFSM) to measure actin dynamics.” Current protocols in cytometry vol. Chapter 2 (2012)). Tracks are shown as red lines. Each displayed area is 8.1 × 8.1 μm^2^ for visual clarity.

## Data availability statement

The data generated for this work are available from the corresponding authors upon reasonable request. The software used for data analysis is previously published and publicly available on GitHub: Colocalization analysis: https://github.com/kjaqaman/conditionalColoc

SMI-FSM analysis: https://github.com/kjaqaman/SMI-FSM

Particle tracking: https://github.com/DanuserLab/u-track

Quantitative fluorescent speckle microscopy: https://github.com/DanuserLab/QFSM

Segmentation: https://github.com/DanuserLab/u-segment

## Acknowledgements

We thank Dr. Tieqiao Zhang for microscopy support. This work was supported by funding from the National Science Foundation (MCB-2114417), the National Institutes of Health/National Institute of General Medical Sciences (R35 GM119619), and the University of Texas Southwestern Endowed Scholars Program to K. Jaqaman. J. Guerrero and J. Vega-Lugo were trainees of the National Institutes of Health Molecular Biophysics Training Grant (5T32GM131963; PI: Dr. Yuh Min Chook).

---

**Figure S1.**
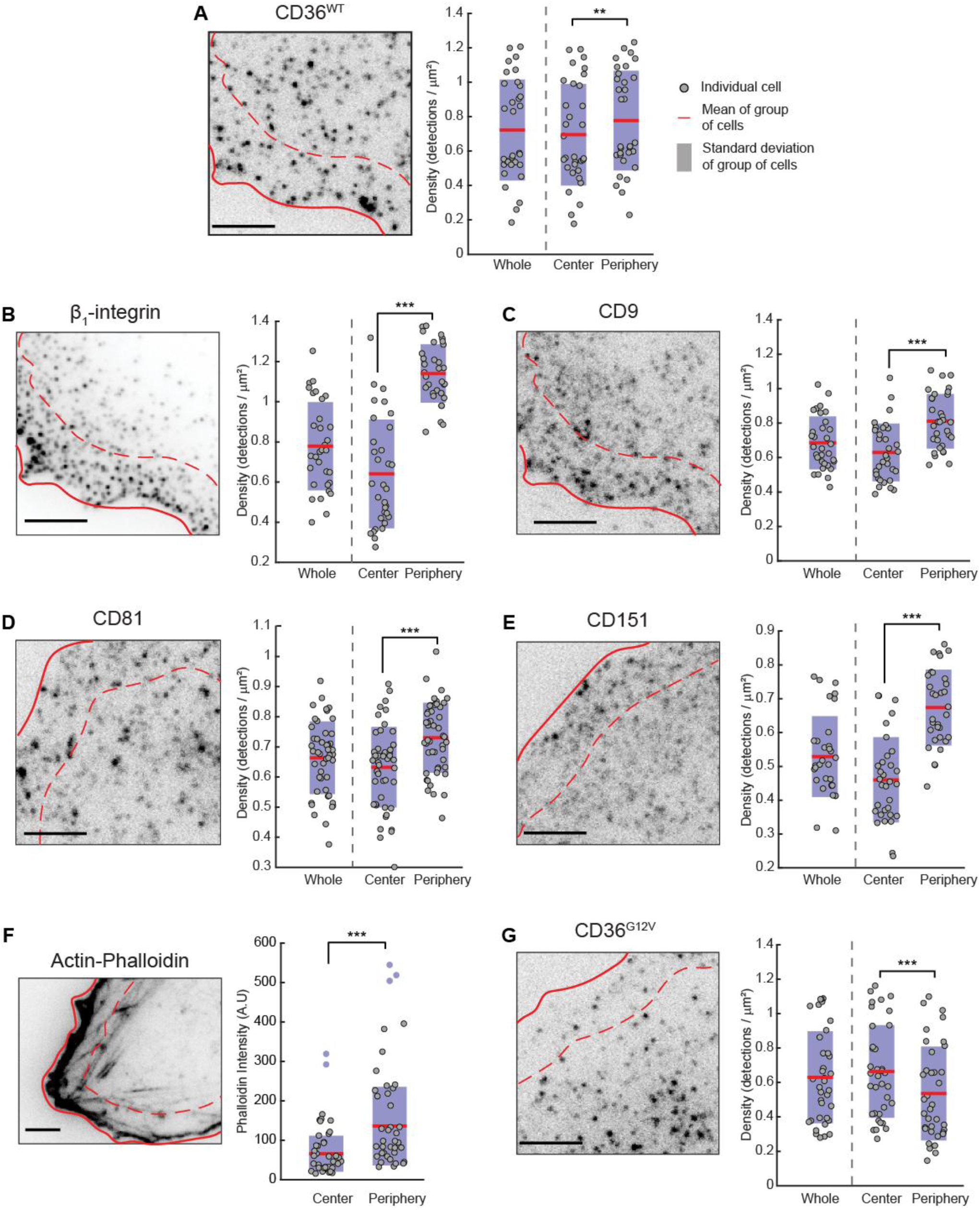
Density distributions of the various imaged molecules and actin in TIME cells. (**A-E** and **G**) Representative fixed-cell images of the indicated surface molecules and quantification of their detection densities in the whole imaged region, as well as in the center and periphery subregions separately. (**F**) Representative fixed-cell image of actin (stained using phalloidin) and quantification of its average intensity in the center and periphery subregions separately. In all images, solid lines show segmented cell edge, dashed lines specify boundary between periphery and center regions. Scale bars, 5 μm. In all plots, circles, red lines, shaded bars, and black asterisks as in Fig 2. Empty circles in F indicate outlier values. See Table 1 for number of cells, number of experimental repeats, and number of objects per channel per ROI used for analysis.

**Figure S2.**
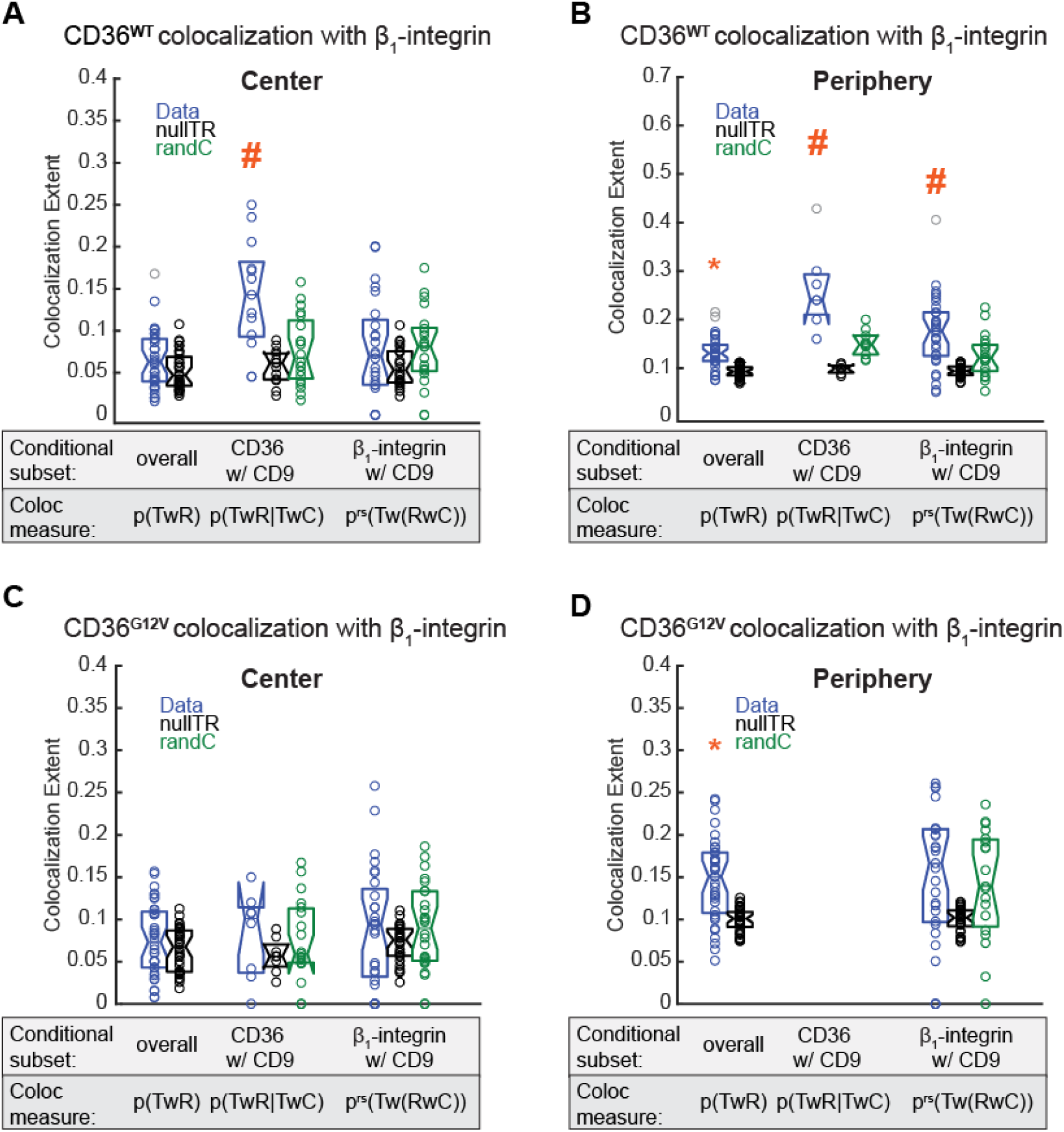
Quantification of CD36 colocalization with β_1_-integrin, conditional on CD9. This figure shows the details of colocalization analysis of the indicated three molecules underlying the p-values presented in Table S1. (**A**) Conditional colocalization measures (blue) and their coincidental counterparts (“nullTR” (black) and “randC” (green)) for target = CD36^WT^, reference = β_1_-integrin, and condition = CD9, in the center region, shown as boxplots. The x-axis lists the colocalization measures and, above them, what molecule subsets they represent. For each box, individual circles indicate individual cell measurements, the central mark is the median, and the edges are the 25^th^ and 75^th^ percentiles. Notch around median indicates the 95% confidence interval of the median. Circles with same color as box are inliers while gray circles are outliers. *: p(TwR) is significantly greater than its coincidental counterpart (“nullTR”) (p ≤ 0.05). #: p(TwR|TwC) or p^rs^(Tw(RwC)) is significantly greater than its coincidental counterparts (both “nullTR” and “randC”) and its corresponding p(TwR) (combined type I error = 0.05 for the 3 tests). (**B**) Same as A, but in the periphery region. (**C, D**), Same as A, B but for CD36^G12V^ in the center (**C**) or periphery (**D**) region. See Table 1 for number of cells, number of experimental repeats, and number of objects per channel per ROI used for analysis.

**Figure S3.**
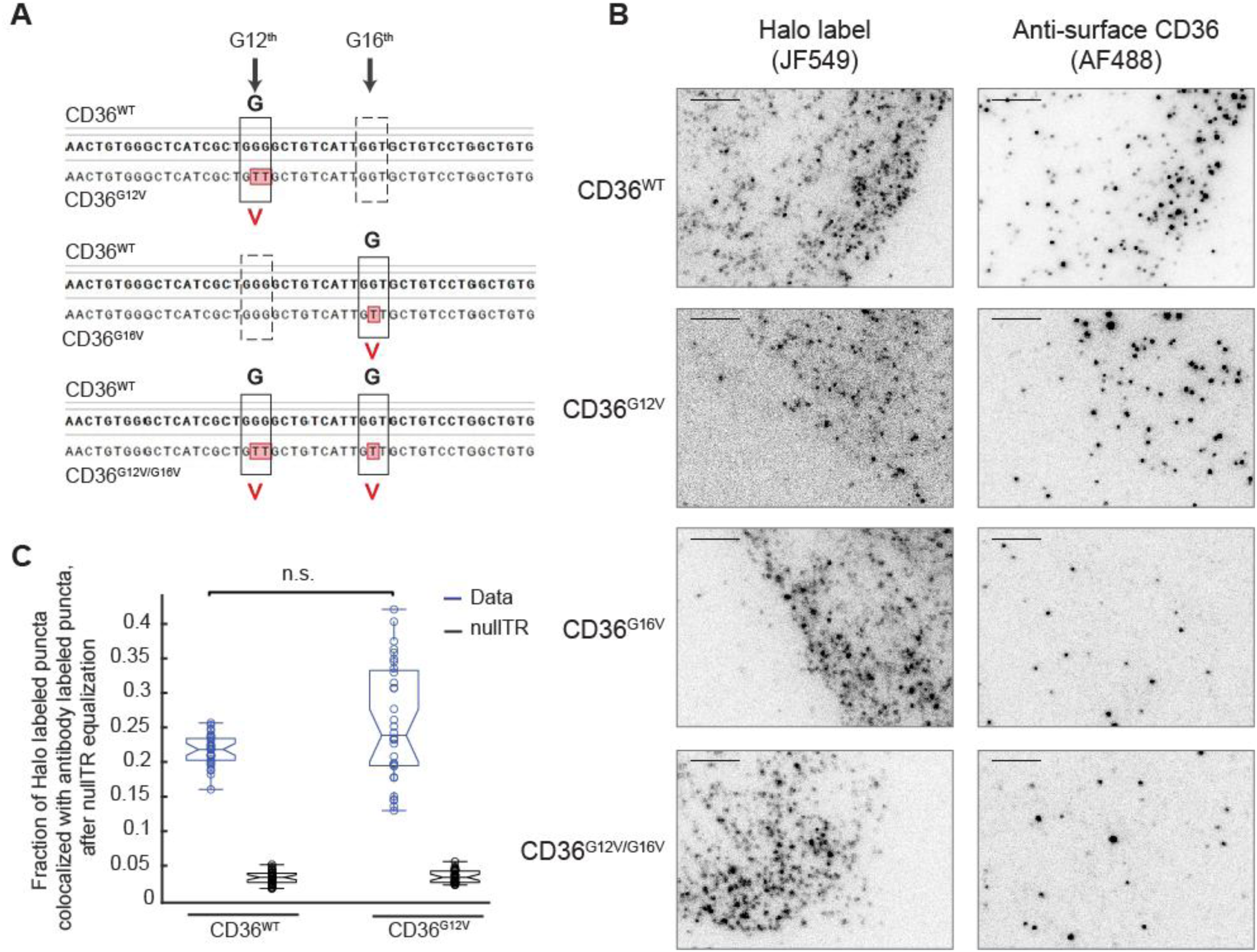
Sequence confirmation and surface expression of different CD36 mutants. (**A**) Sequence of wildtype (bold) compared to CD36^G12V^, CD36^G16V^, and CD36^G12V/G16V^. The change in codon of glycine to valine is shown in black solid box and the nucleotides are highlighted in red color. (**B**) Two-color TIRFM images of HaloTag-fused WT or mutated CD36 (CD36^G12V^, CD36^G16V^, or CD36^G12V/G16V^) expressed in TIME cells, labeled using JF549-conjugated Halo ligand to assess total expression (first column) and an anti-CD36 antibody that targets its ectodomain to assess surface expression (second column). Scale bars, 5 μm. (**C**) To assess the fraction of CD36^WT^ and CD36^G12V^ expressed on the cell surface, the fraction of Halo-labeled puncta (labeling CD36 regardless of its location) colocalized with antibody-labeled puncta (labeling only CD36 on the cell surface) was calculated. The plot shows these fractions after compensating for the slightly different overall expression levels of WT and mutant CD36, achieved by nullTR equalization. NullTR equalization = multiplying the data and nullTR values of one condition (CD36^WT^ in this case) with a factor to make its nullTR have the same median as the nullTR of the condition it is compared to (CD36^G12V^ in this case). The idea is that this compensates for different number of objects in the two conditions, as nullTR reflects the number of objects used in colocalization analysis (Vega-Lugo et al, JCB 2022), thus allowing the direct comparison of the colocalization fractions between the two conditions. Plot details as in Fig. S2. N = 32 cells for each of CD36^WT^ and CD36^G12V^, from 4 experimental repeats.

**Figure S4.**
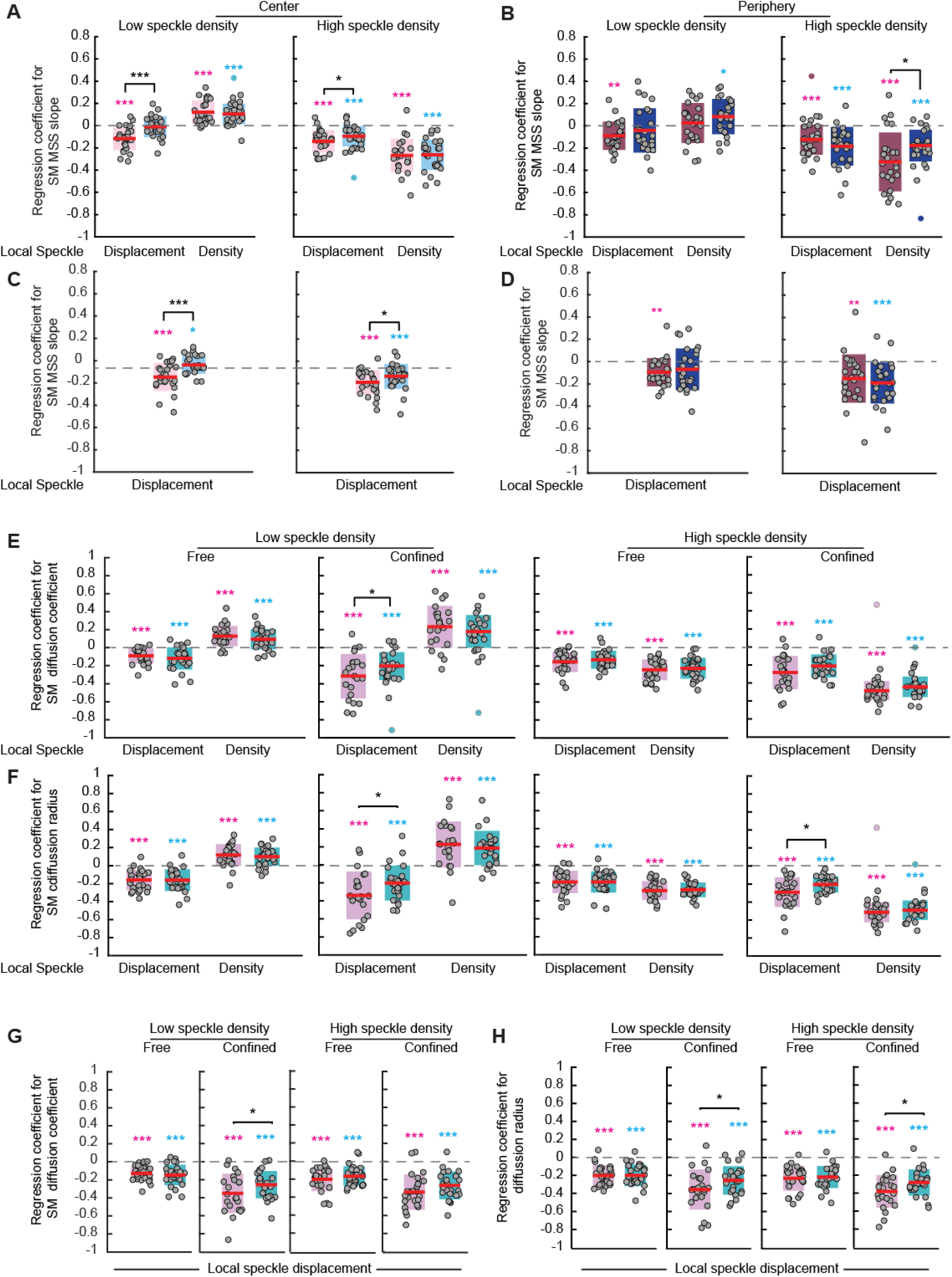
Regression coefficients from MVRG analysis on local CA speckle displacement and density, and from MVRG analysis on local CA speckle displacement alone. (**A, B**) Regression coefficients of CD36^WT^ and CD36^G12V^ MSS slope in the center (**A**) and periphery (**B**) regions from MVRG analysis on speckle displacement and density. This is the same MVRG analysis in Fig. 2A, B, but showing both regression coefficient components here (while Fig. 2A, B show only the displacement component). (**C, D**) Regression coefficients of CD36^WT^ and CD36^G12V^ MSS slope in the center (**C**) and periphery (**D**) regions from MVRG analysis on speckle displacement alone. (**E, F**) Regression coefficients of CD36^WT^ and CD36^G12V^ diffusion coefficient (**E**) and diffusion radius (**F**) on speckle displacement and density. This is the same MVRG analysis in Fig. 4A, B, but showing both regression coefficient components here (while Fig. 4A, B shows only the displacement component). (**G, H**) Regression coefficients of CD36^WT^ and CD36^G12V^ diffusion coefficient (**G**) and diffusion radius (**H**) from MVRG analysis on speckle displacement alone. All figure details as in Fig. 2 and Fig. 4. Comparing A vs. C, B vs. D, E vs. G and F vs. H shows very similar regression coefficients on displacement whether MVRG is performed on displacement and density or displacement alone. At the same time, including the density in the MVRG analysis yields somewhat cleaner results for the displacement regression coefficients (e.g., the analysis at high local CA speckle density in B vs. D). See Table 2 for number of cells, number of experimental repeats, and number of SM tracklets and speckles per ROI used for analysis.

**Table S1.**
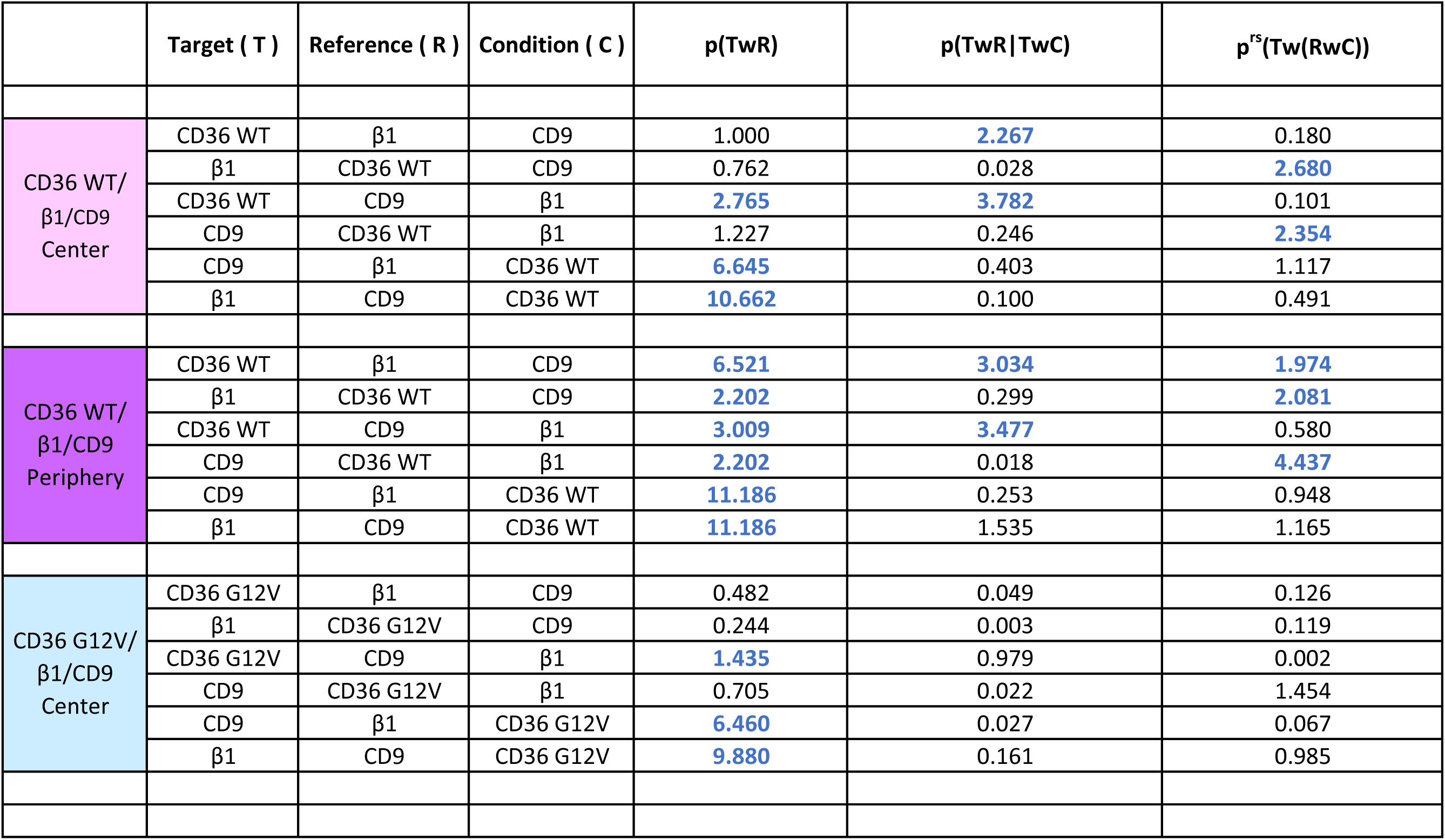
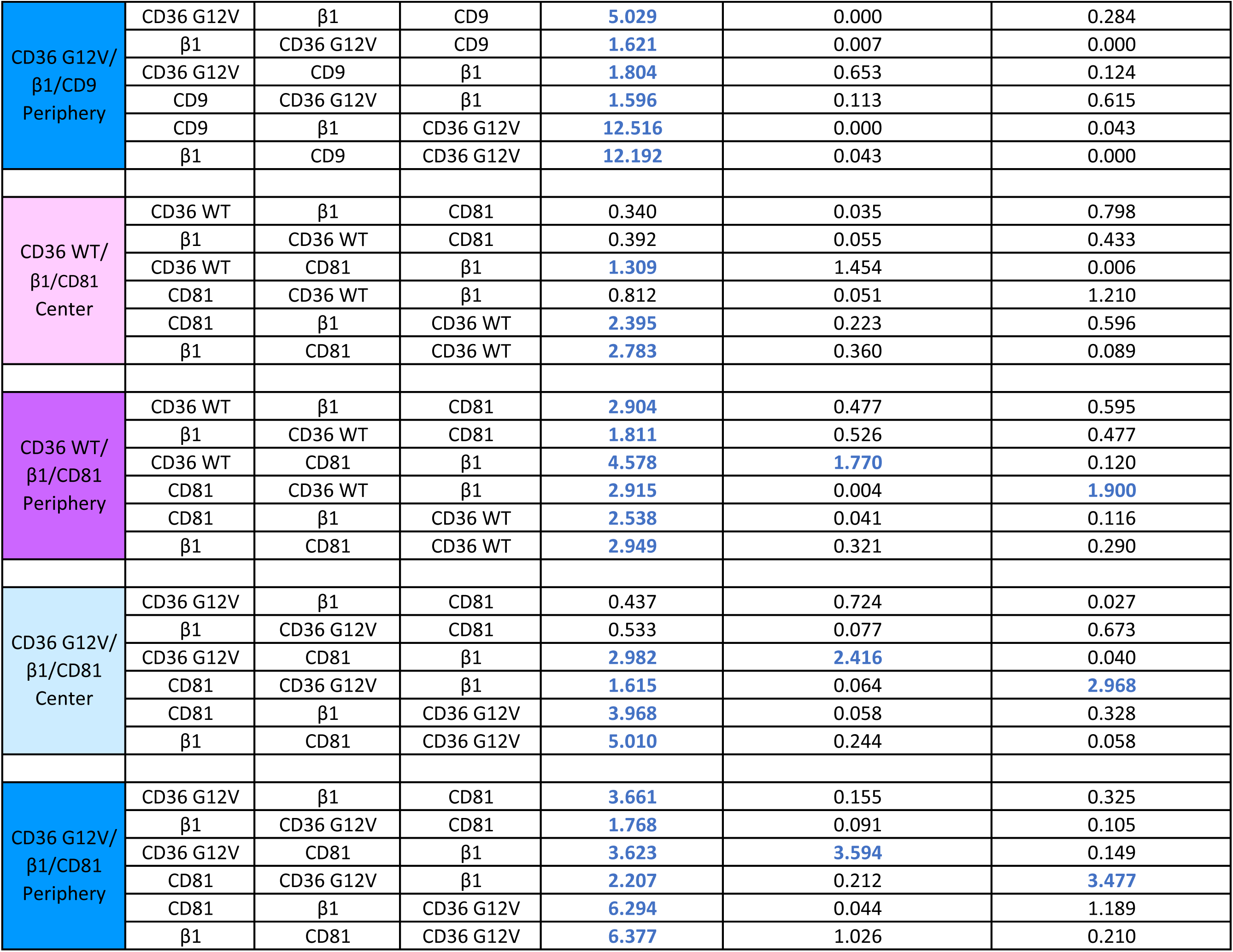
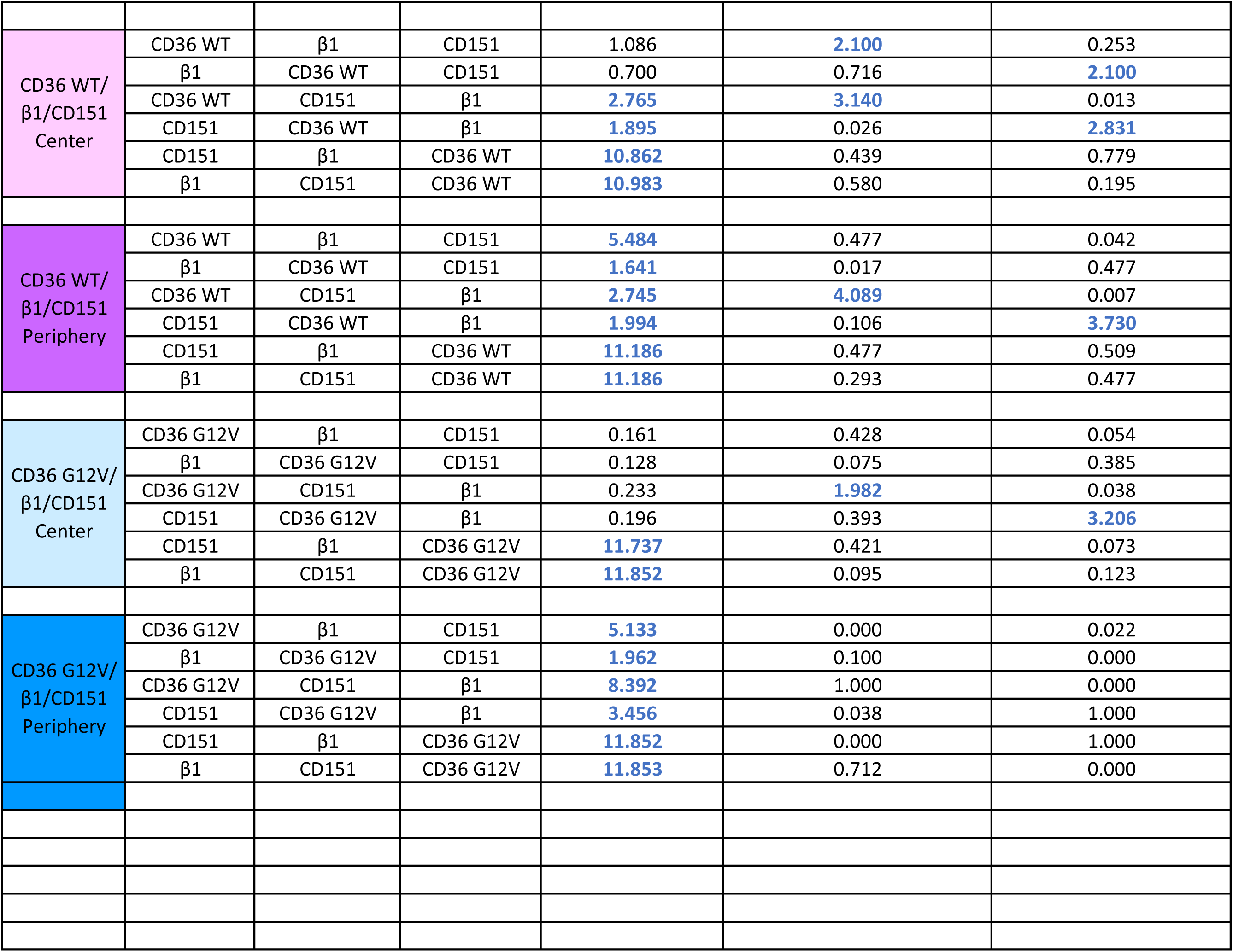
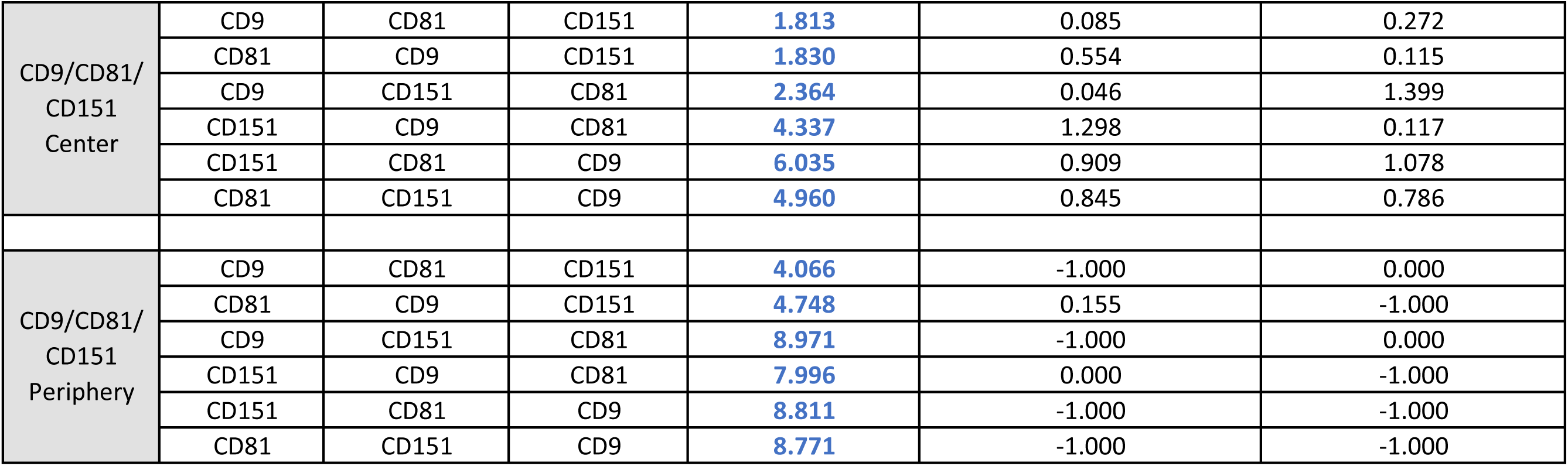
−log10(significance p-values) for the indicated colocalization and conditional colocalization measures in the center and periphery ROIs, for the listed datasets and permutations of molecules into target, reference and condition. These −log10(significance p-values) were used to generate the colocalization networks shown in Fig. 1. In the table, −log10(p-values) ≥ the significance threshold, i.e. −log10(0.05) = 1.301 for p(TwR) and −log10(0.017) = 1.770 for p(TwR|TwC) and p^rs^(Tw(RwC)), are pointed out in blue. In the CD9/CD81/CD151 periphery ROI analysis, −1 indicates not enough datapoints in any analyzed ROI to calculate the indicated conditional colocalization measure.

| | Target ( T ) | Reference ( R ) | Condition ( C ) | $p(\text{TwR})$ | $p(\text{TwR}\vert \text{TwC})$ | $p^{\text{rs}}(\text{Tw}(\text{RwC}))$ |
| --- | --- | --- | --- | --- | --- | --- |
| CD36 WT/<br>$\beta 1$/CD9<br>Center | CD36 WT | $\beta 1$ | CD9 | 1.000 | 2.267 | 0.180 |
| | $\beta 1$ | CD36 WT | CD9 | 0.762 | 0.028 | 2.680 |
| | CD36 WT | CD9 | $\beta 1$ | 2.765 | 3.782 | 0.101 |
| | CD9 | CD36 WT | $\beta 1$ | 1.227 | 0.246 | 2.354 |
| | CD9 | $\beta 1$ | CD36 WT | 6.645 | 0.403 | 1.117 |
| | $\beta 1$ | CD9 | CD36 WT | 10.662 | 0.100 | 0.491 |
| CD36 WT/<br>$\beta 1$/CD9<br>Periphery | CD36 WT | $\beta 1$ | CD9 | 6.521 | 3.034 | 1.974 |
| | $\beta 1$ | CD36 WT | CD9 | 2.202 | 0.299 | 2.081 |
| | CD36 WT | CD9 | $\beta 1$ | 3.009 | 3.477 | 0.580 |
| | CD9 | CD36 WT | $\beta 1$ | 2.202 | 0.018 | 4.437 |
| | CD9 | $\beta 1$ | CD36 WT | 11.186 | 0.253 | 0.948 |
| | $\beta 1$ | CD9 | CD36 WT | 11.186 | 1.535 | 1.165 |
| CD36 G12V/<br>$\beta 1$/CD9<br>Center | CD36 G12V | $\beta 1$ | CD9 | 0.482 | 0.049 | 0.126 |
| | $\beta 1$ | CD36 G12V | CD9 | 0.244 | 0.003 | 0.119 |
| | CD36 G12V | CD9 | $\beta 1$ | 1.435 | 0.979 | 0.002 |
| | CD9 | CD36 G12V | $\beta 1$ | 0.705 | 0.022 | 1.454 |
| | CD9 | $\beta 1$ | CD36 G12V | 6.460 | 0.027 | 0.067 |
| | $\beta 1$ | CD9 | CD36 G12V | 9.880 | 0.161 | 0.985 |
| CD36 G12V/<br>β1/CD9<br>Periphery | CD36 G12V | β1 | CD9 | 5.029 | 0.000 | 0.284 |
|  | β1 | CD36 G12V | CD9 | 1.621 | 0.007 | 0.000 |
|  | CD36 G12V | CD9 | β1 | 1.804 | 0.653 | 0.124 |
|  | CD9 | CD36 G12V | β1 | 1.596 | 0.113 | 0.615 |
|  | CD9 | β1 | CD36 G12V | 12.516 | 0.000 | 0.043 |
|  | β1 | CD9 | CD36 G12V | 12.192 | 0.043 | 0.000 |
| CD36 WT/<br>β1/CD81<br>Center | CD36 WT | β1 | CD81 | 0.340 | 0.035 | 0.798 |
|  | β1 | CD36 WT | CD81 | 0.392 | 0.055 | 0.433 |
|  | CD36 WT | CD81 | β1 | 1.309 | 1.454 | 0.006 |
|  | CD81 | CD36 WT | β1 | 0.812 | 0.051 | 1.210 |
|  | CD81 | β1 | CD36 WT | 2.395 | 0.223 | 0.596 |
|  | β1 | CD81 | CD36 WT | 2.783 | 0.360 | 0.089 |
| CD36 WT/<br>β1/CD81<br>Periphery | CD36 WT | β1 | CD81 | 2.904 | 0.477 | 0.595 |
|  | β1 | CD36 WT | CD81 | 1.811 | 0.526 | 0.477 |
|  | CD36 WT | CD81 | β1 | 4.578 | 1.770 | 0.120 |
|  | CD81 | CD36 WT | β1 | 2.915 | 0.004 | 1.900 |
|  | CD81 | β1 | CD36 WT | 2.538 | 0.041 | 0.116 |
|  | β1 | CD81 | CD36 WT | 2.949 | 0.321 | 0.290 |
| CD36 G12V/<br>β1/CD81<br>Center | CD36 G12V | β1 | CD81 | 0.437 | 0.724 | 0.027 |
|  | β1 | CD36 G12V | CD81 | 0.533 | 0.077 | 0.673 |
|  | CD36 G12V | CD81 | β1 | 2.982 | 2.416 | 0.040 |
|  | CD81 | CD36 G12V | β1 | 1.615 | 0.064 | 2.968 |
|  | CD81 | β1 | CD36 G12V | 3.968 | 0.058 | 0.328 |
|  | β1 | CD81 | CD36 G12V | 5.010 | 0.244 | 0.058 |
| CD36 G12V/<br>β1/CD81<br>Periphery | CD36 G12V | β1 | CD81 | 3.661 | 0.155 | 0.325 |
|  | β1 | CD36 G12V | CD81 | 1.768 | 0.091 | 0.105 |
|  | CD36 G12V | CD81 | β1 | 3.623 | 3.594 | 0.149 |
|  | CD81 | CD36 G12V | β1 | 2.207 | 0.212 | 3.477 |
|  | CD81 | β1 | CD36 G12V | 6.294 | 0.044 | 1.189 |
|  | β1 | CD81 | CD36 G12V | 6.377 | 1.026 | 0.210 |
| CD36 WT/<br>β1/CD151<br>Center | CD36 WT | β1 | CD151 | 1.086 | 2.100 | 0.253 |
| | β1 | CD36 WT | CD151 | 0.700 | 0.716 | 2.100 |
| | CD36 WT | CD151 | β1 | 2.765 | 3.140 | 0.013 |
| | CD151 | CD36 WT | β1 | 1.895 | 0.026 | 2.831 |
| | CD151 | β1 | CD36 WT | 10.862 | 0.439 | 0.779 |
| | β1 | CD151 | CD36 WT | 10.983 | 0.580 | 0.195 |
| CD36 WT/<br>β1/CD151<br>Periphery | CD36 WT | β1 | CD151 | 5.484 | 0.477 | 0.042 |
| | β1 | CD36 WT | CD151 | 1.641 | 0.017 | 0.477 |
| | CD36 WT | CD151 | β1 | 2.745 | 4.089 | 0.007 |
| | CD151 | CD36 WT | β1 | 1.994 | 0.106 | 3.730 |
| | CD151 | β1 | CD36 WT | 11.186 | 0.477 | 0.509 |
| | β1 | CD151 | CD36 WT | 11.186 | 0.293 | 0.477 |
| CD36 G12V/<br>β1/CD151<br>Center | CD36 G12V | β1 | CD151 | 0.161 | 0.428 | 0.054 |
| | β1 | CD36 G12V | CD151 | 0.128 | 0.075 | 0.385 |
| | CD36 G12V | CD151 | β1 | 0.233 | 1.982 | 0.038 |
| | CD151 | CD36 G12V | β1 | 0.196 | 0.393 | 3.206 |
| | CD151 | β1 | CD36 G12V | 11.737 | 0.421 | 0.073 |
| | β1 | CD151 | CD36 G12V | 11.852 | 0.095 | 0.123 |
| CD36 G12V/<br>β1/CD151<br>Periphery | CD36 G12V | β1 | CD151 | 5.133 | 0.000 | 0.022 |
| | β1 | CD36 G12V | CD151 | 1.962 | 0.100 | 0.000 |
| | CD36 G12V | CD151 | β1 | 8.392 | 1.000 | 0.000 |
| | CD151 | CD36 G12V | β1 | 3.456 | 0.038 | 1.000 |
| | CD151 | β1 | CD36 G12V | 11.852 | 0.000 | 1.000 |
| | β1 | CD151 | CD36 G12V | 11.853 | 0.712 | 0.000 |
| CD9/CD81/<br>CD151<br>Center | CD9 | CD81 | CD151 | 1.813 | 0.085 | 0.272 |
|  | CD81 | CD9 | CD151 | 1.830 | 0.554 | 0.115 |
|  | CD9 | CD151 | CD81 | 2.364 | 0.046 | 1.399 |
|  | CD151 | CD9 | CD81 | 4.337 | 1.298 | 0.117 |
|  | CD151 | CD81 | CD9 | 6.035 | 0.909 | 1.078 |
|  | CD81 | CD151 | CD9 | 4.960 | 0.845 | 0.786 |
| CD9/CD81/<br>CD151<br>Periphery | CD9 | CD81 | CD151 | 4.066 | -1.000 | 0.000 |
|  | CD81 | CD9 | CD151 | 4.748 | 0.155 | -1.000 |
|  | CD9 | CD151 | CD81 | 8.971 | -1.000 | 0.000 |
|  | CD151 | CD9 | CD81 | 7.996 | 0.000 | -1.000 |
|  | CD151 | CD81 | CD9 | 8.811 | -1.000 | -1.000 |
|  | CD81 | CD151 | CD9 | 8.771 | -1.000 | -1.000 |

